# Sphingolipids are involved in *Pieris brassicae* egg-induced cell death in *Arabidopsis thaliana*

**DOI:** 10.1101/2021.07.09.451813

**Authors:** Raphaël Groux, Laetitia Fouillen, Sébastien Mongrand, Philippe Reymond

**Affiliations:** Department of Plant Molecular Biology, University of Lausanne, CH-1015 Lausanne, Switzerland; Laboratoire de Biogénèse Membranaire, CNRS, UMR 5200, University of Bordeaux, F-33140 Villenave d’Ornon, France

## Abstract

In *Brassicaceae*, hypersensitive-like (HR-like) cell death is a central component of direct defenses triggered against eggs of the large white butterfly *Pieris brassicae*. The signaling pathway leading to HR-like in Arabidopsis is mainly dependent on salicylic acid (SA) accumulation, but downstream components are unclear. Here, we found that treatment with *P. brassicae* egg extract (EE) trigger changes in expression of sphingolipid metabolism genes in Arabidopsis and *Brassica nigra*. Disruption of ceramide synthase activity led to a significant decrease of EE-induced HR-like whereas SA signaling and reactive oxygen species levels were unchanged, suggesting that ceramides are downstream activators of HR-like. Sphingolipid quantifications showed that ceramides with C16:0 side-chains accumulated in both species, and this response was independent on SA accumulation. Finally, we provide genetic evidence that the modification of fatty acyl chains of sphingolipids modulates HR-like. Altogether, these results show that sphingolipids play a key and specific role during insect egg-triggered HR-like.

## Introduction

Programmed cell death (PCD) plays an essential role in plants. It is part of development by promoting cell and tissue differentiation but results also from immune defense system activation (Reape and McCabe, 2010; Coll et al., 2011; Huysmans et al., 2017). The best studied form of pathogen-triggered PCD is termed the hypersensitive response (HR), a spectacular response triggered upon recognition of adapted pathogens by resistance proteins that leads to macroscopic cell death, induction of defense gene expression and pathogen resistance (Balint-Kurti, 2019). A meta-analysis of PCD-inducing conditions revealed that transcriptomic signatures of developmental PCD and pathogen-triggered PCD are largely distinct (Olvera-Carrillo et al., 2015), suggesting that they are under different genetic regulation. More specifically, pathogen-triggered PCD is dependent on salicylic acid (SA) accumulation and signaling (Coll et al., 2011; Huysmans et al., 2017; Balint-Kurti, 2019). In addition to immunity to pathogens, it was reported that hypersensitivity may also function as a defense strategy against insect herbivores (Fernandes, 1990; Stuart, 2015). In particular, plants from the *Brassicales, Solanales* and *Fabales* were shown to induce localized cell death in response to oviposition by insects (Shapiro and DeVay, 1987; Balbyshev and Lorenzen, 1997; Garza et al., 2001; Little et al., 2007; Petzold-Maxwell et al., 2011; Fatouros et al., 2016; Geuss et al., 2017; Griese et al., 2021), a process called HR-like (Reymond, 2013; Fatouros et al., 2014). As a consequence, direct defense induction correlates with decreased egg survival and/or increased egg parasitism (Shapiro and DeVay, 1987; Balbyshev and Lorenzen, 1997; Fatouros et al., 2014; Fatouros et al., 2016; Geuss et al., 2017; Griese et al., 2017; Griese et al., 2021). Like pathogen-triggered HR, egg-induced HR-like responses are associated with an accumulation of reactive oxygen species (ROS) and SA, and defense gene expression (Little et al., 2007; Hilfiker et al., 2014; Geuss et al., 2017; Bonnet et al., 2017). Studies in *Arabidopsis thaliana* reported that the signaling cascade involved in the response to eggs of the Large White Butterfly *Pieris brassicae* is similar to pathogen-triggered immunity (PTI) (Gouhier-Darimont et al. 2013). Notably, the induction of cell death was dependent on SA accumulation and signaling. The exact cause of the decreased egg survival associated with HR-like is not known, but data from *Brassica nigra* suggest that it could be due to water removal at the oviposition site (Griese et al., 2017), consistent with low water potential observed in tissues undergoing HR (Wright and Beattie, 2004). In addition, exposure to ROS at the oviposition site was shown to increase egg mortality (Geuss et al., 2017). These data thus suggest that HR-like at oviposition sites may constitute an efficient defense strategy against insect eggs.

As it could decrease insect pressure before damage occurs, the introgression of egg-killing traits in cultivated crop species is desirable (Fatouros et al., 2016) and has been successfully reported in *Oryza sativa* (Suzuki et al., 1996; Yamasaki et al., 2003; Yang et al., 2014). Despite this achievement, this strategy is still mostly overlooked as this response is poorly understood at the molecular level (Reymond, 2013; Fatouros et al., 2016). The use of Arabidopsis as a model plant to explore the genetic basis of the response to *P. brassicae* eggs has so far successfully identified PTI components as regulators of egg-induced HR-like and showed that activation of cell surface receptor-like kinases LecRK-I.1 and LecRK-I.8 is an early step of egg-induced responses (Gouhier-Darimont et al., 2013; Gouhier-Darimont et al., 2019). Moreover, phosphatidylcholines derived from *P. brassicae* egg extract were recently shown to induce defense responses and cell death (Stahl et al., 2020). However, the identity of cell-death inducing factors downstream of SA is unknown.

In contrast to animals, plants lack certain central components of PCD pathways, such as caspases (Coll et al., 2011; Salvesen et al., 2015), but instead rely on a variety of other proteases that fulfill similar functions (Salguero-Linares and Coll, 2019). The identification and characterization of lesion mimic mutants, which display spontaneous cell death along with elevated defenses, has largely contributed to shed light on processes involved in PCD (Bruggeman et al., 2015). In particular, several lesion mimic mutants were found to function in sphingolipid metabolism. The involvement of sphingolipids in PCD induction in animals is well described (Young et al., 2013), and their function is conserved in plants (Townley et al., 2005; Huby et al., 2019). Sphingolipids differ from glycerolipids as they consist of a sphingoid long-chain base (LCB) linked *via* the amide bond to one fatty acid (FA) moiety (Ali et al., 2018). LCB backbones can be further modified through an α-hydroxylation or a desaturation. These molecules, called ceramides (Cer), can be further modified by the attachment of a polar head group consisting of a glucose or a glycosyl inositol phosphoryl moiety, leading to the formation of complex sphingolipids such as GluCer (glucosylceramides) or GIPC (glycosyl inositol phospho ceramides), respectively. In plants, the large majority of identified sphingolipids are complex (Markham et al., 2013; Gronnier et al, 2016; Carmon-Salazar et al., 2021), whereas LCB and Cer are low abundant. Interestingly, both free LCB and Cer have been shown to induce PCD when exogenously applied to plants (Liang et al., 2003; Shi et al., 2007; Lachaud et al., 2011; Saucedo-García et al., 2011). Additionally, several fungal toxins such as Fumonisin B1 were shown to cause cell death through an accumulation of free LCB by inhibiting ceramide synthases (Berkey et al., 2012). While the mechanisms involved downstream of LCB/Cer are not clear, the modification of sphingolipid levels in the context of immune responses was shown to affect pathogen resistance (Ternes et al., 2011; Magnin-Robert et al., 2015; Wu et al., 2015). Interestingly, one study found a role for sphingolipid metabolism in resistance against insects. Expression of *OsLCB2*, encoding a serine palmitoyl transferase involved in the first step of LCB biosynthesis, was found to be induced by brown planthopper infestation and overexpression of *OsLCB2* in Arabidopsis triggered LCB accumulation, SA-dependent gene expression and resistance to aphids (Begum et al., 2016).

Here we report that eggs of *P. brassicae* alter the expression of sphingolipid metabolism genes and trigger an accumulation of ceramides in both Arabidopsis and *B. nigra.* Furthermore, we show that HR-like induction is affected in different ceramide synthase and FA hydroxylases mutants, whereas ROS and SA levels are not impaired in the mutants. Altogether, these data indicate that sphingolipids play a key role in the execution of egg-induced cell death.

## Results

### *P. brassicae* eggs induce biotic cell death markers

Different types of PCD exist in plants and a meta-analysis of publicly available transcriptomic data previously enabled the identification of marker genes for different types of cell death: biotic, osmotic, developmental and genotoxic (Olvera-Carrillo et al., 2015). We previously published transcriptomic data from Arabidopsis plants subjected to natural oviposition (Little et al., 2007) and used these expression profiles to explore the molecular signatures associated with egg-induced HR-like. We extracted expression data for the different PCD marker genes described in Olvera-Carillo et al. (2015) 24 h, 48 h and 72 h after egg deposition by *P. brassicae*. Interestingly, marker genes for biotic cell death were found to be highly induced after egg deposition, while markers for other types of PCD were weakly responsive (Figure 1).

**Figure 1.**
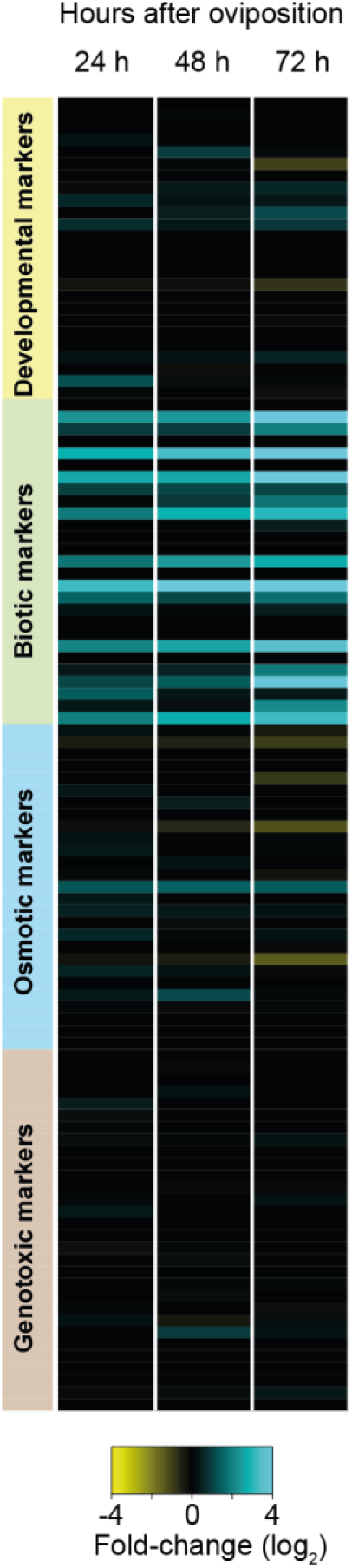
Hypersensitive-like cell death following *P. brassicae* oviposition induces markers of biotic PCD. Expression of marker genes for biotic-related PCD in Arabidopsis plants following oviposition by *P. brassicae* butterflies for 24, 48 or 72 h. Marker genes were described in Olvera-Carrillo et al. (2015) and expression data were extracted from a previously published microarray study by Little et al. (2007).

### Expression of lipid metabolism genes is altered in response to *P. brassicae* oviposition

Lipid metabolism is central in plant development and some sectors have been shown to be involved in PCD during immunity (Siebers et al., 2016; Lim et al., 2017). We explored the potential involvement of lipid metabolism during egg-induced responses. Using transcriptome data from *P. brassicae* oviposition on Arabidopsis after 24, 48 and 72 h (Little et al., 2007), we extracted expression ratios for genes related to lipid metabolism (AraLip database; http://aralip.plantbiology.msu.edu/). Only genes whose expression was significantly different (ratio ≥|1.5|, adj P value <0.05) at least at one time point were selected. This analysis led to a list of 136 genes (out of 765 in the AraLip database) representative of all major lipid pathways (Figure 2A). Data clustering showed that genes were either up-or downregulated over time, displaying a very sharp regulation process. Interestingly, genes involved in processes such as FA synthesis, elongation or phospholipid synthesis were mostly downregulated while genes in sphingolipid biosynthesis, TAG degradation, suberin and oxylipin biosynthesis were mainly upregulated (Figure 2B). Notably, both oxylipins and sphingolipids have previously been involved in the regulation of cell death (Siebers et al., 2016; Lim et al., 2017; Huby et al., 2019), hinting to a potential implication during egg-induced responses.

**Figure 2.**
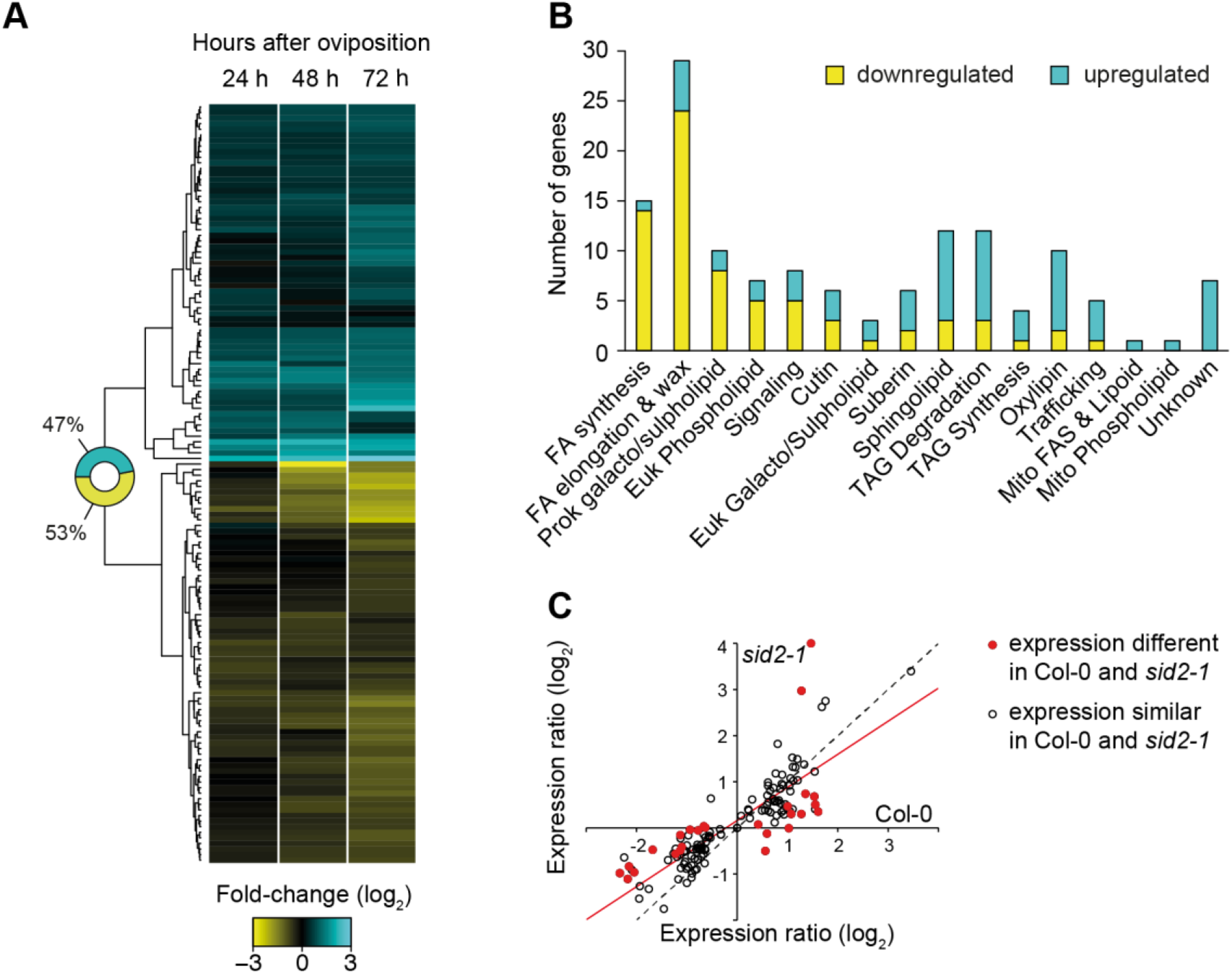
Transcriptomic alterations in lipid metabolism after insect egg deposition. A, Heatmap showing expression of genes involved in lipid metabolism after oviposition by *P. brassicae* on Arabidopsis plants. Microarray data were taken from Little et al. (2007) and a list of genes specifically involved in lipid metabolism was obtained from the AraLip database. Only genes that were differentially regulated between control and treated plants (Fold-change ≥ |1.5|, adj *P*<0.05) in at least one time-point are shown. B, Number of gene up- or down-regulated in each metabolic categories defined on the AraLip database. C, Expression of lipid metabolism genes in Col-0 and *sid2-1* mutant plants three days after egg deposition. Expression ratios from Col-0 are plotted against expression ratios from *sid2-1*. Each circle represents one gene that is induced by eggs in Col-0. Filled circles are genes whose expression was significantly different in *sid2-1*; open circles are genes whose expression was not different between Col-0 and *sid2-1*. The dotted line indicates perfect correspondence in expression ratios between Col-0 and *sid2-1* while the red line represents a regression analysis of the dataset (y = 0.72x + 0.16, R² = 0.71).

Biotic PCD is typically induced upon recognition of pathogens and this process is regulated by SA (Radojičić et al., 2018). In addition, we previously showed that *P. brassicae* eggs trigger responses that require the SA pathway (Bruessow et al., 2010; Gouhier-Darimont et al., 2013). We thus examined whether transcriptional alterations of lipid metabolism genes were dependent on SA accumulation. Looking at expression of lipid metabolism genes in the published oviposition transcriptome data with the SA biosynthesis mutant *sid2-1* (Little et al., 2007), we found only a few genes that displayed significantly altered transcript levels after oviposition on *sid2-1* compared to Col-0 (Figure 2C), indicating that the transcriptional reprogramming of lipid metabolism is mainly independent from SA accumulation. However, linear fitting of both datasets shows that, overall, changes in gene expression were lower in *sid2-1* (as seen by regression line closer to the Col-0 axis), suggesting a partial contribution of SA signaling to this response (Figure 2C).

### HR-like induction is independent of MYB30 and oxylipin synthesis

We further explored the possibility that lipid metabolism may play a role in HR-like induction upon insect egg perception. MYB30 was previously shown to regulate pathogen-induced HR through the transcriptional regulation of VLCFA biosynthesis and accumulation (Raffaele et al., 2008), providing an interesting link between lipid metabolism and cell death induction. Because most VLCFA are found in sphingolipids and cuticular waxes (De Bigault Du Granrut and Cacas, 2016), the authors concluded that MYB30 induces cell death by promoting substrate accumulation for sphingolipid synthesis (Raffaele et al., 2008; De Bigault Du Granrut and Cacas, 2016). As *MYB30* expression was transiently induced before cell death onset, we measured the expression of both *MYB30* and *FATB*, one of its target gene (Raffaele et al., 2008), during the first 24 h after *P. brassicae* crude egg extract (EE) treatment. Mutant plants were treated with EE, which mimics responses induced by natural oviposition (Little et al., 2007; Bruessow et al., 2010; Gouhier-Darimont et al., 2013; Hilfiker et al., 2014; Stahl et al., 2020). However, neither of these genes was induced upon treatment and *FATB* expression was even repressed over time (Supplemental Figure S1). In addition, previous microarray data showed that *MYB30* is repressed later during the EE response, along with other MYB30-regulated genes (Little et al., 2007). Finally, EE-triggered cell death, quantified by trypan blue staining (Gouhier-Darimont et al., 2013), was not altered in *myb30*, indicating that this gene is not involved in the induction of HR-like (Figure 3A). These data are in agreement with the observed repression of FA synthesis/elongation genes (Figure 2A, B).

**Figure 3.**
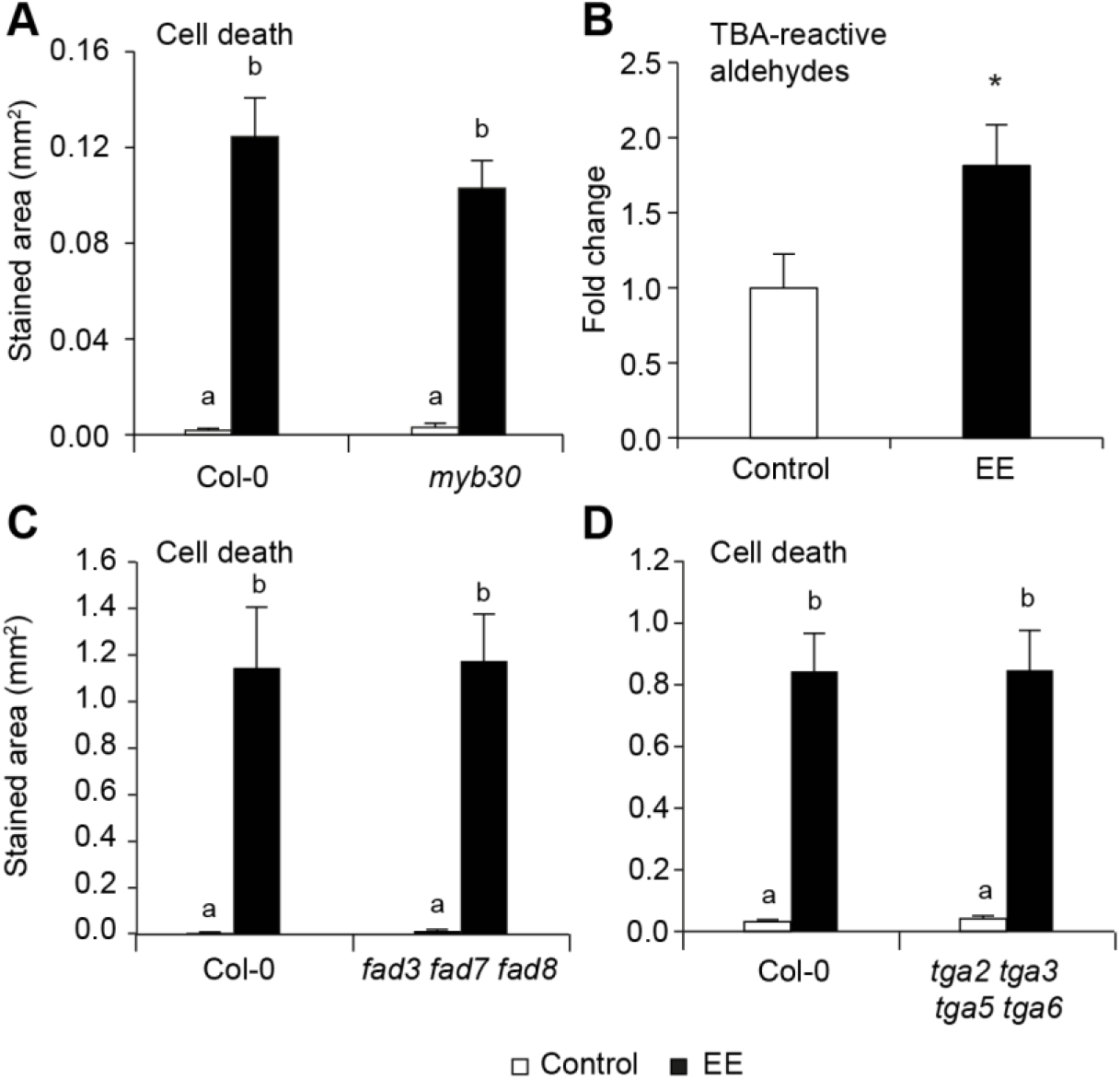
EE-induced cell death is independent from MYB30 and oxylipins. A, C, D, Cell death after three days of EE treatments in different mutants. For each genotype, a total of 8-12 leaves from 4 to 6 plants were treated with 2 μL of EE and cell death was quantitated by trypan blue staining. Untreated leaves were used as controls. All experiments were repeated twice with similar results. Different letters indicate significant differences at *P*<0.05 (ANOVA, followed by Tukey’s HSD for multiple comparisons). B, Relative nonenzymatic lipid peroxidation levels were measured by the quantification of TBA reactive aldehydes in Col-0 plants treated with EE for three days. Data represent means ± SE of three independent experiments (n = 6 leaf discs/experiment). Asterisks denote statistically significant differences (Welch *t-* test, *, *P*<0.05).

Lipid peroxidation plays a crucial role in the regulation of cell death through the production of oxylipins (García-Marcos et al., 2013; Siebers et al., 2016), and it was reported that important oxylipin production occurs upon induction of HR by bacterial pathogens (Andersson et al., 2006). This process occurs upon enzymatic or non-enzymatic polyunsaturated FA oxidation, and one of the best known oxylipin is jasmonic acid. We assessed the level of lipid peroxidation after 3 days of EE treatment by using the thiobarbituric acid assay, which gives an indirect measure of lipid peroxidation through the detection of its byproducts (Stahl et al., 2019). Interestingly, we observed that EE caused an increase in the level of lipid peroxidation in wild-type plants (Figure 3B), a clear argument that oxylipins are indeed produced in response to eggs. We then genetically assessed whether lipid peroxidation was necessary for cell death induction by using the FA desaturase *fad3fad7fad8* triple mutant, which lacks tri-unsaturated FAs from which most oxylipins are derived (McConn and Browse, 1996; Weber et al., 2004). Trypan blue staining following EE treatment did not reveal any difference in the ability of *fad3fad7fad8* mutant plants to induce cell death, demonstrating that this process is independent from trienoic FAs and oxylipin production. Furthermore, TGA transcription factors TGA2, TGA5 and TGA6 were shown to transduce responses downstream of oxylipins such as OPDA and phytoprostanes (Mueller et al., 2008; Stotz et al., 2013). Consistent with our previous results, the quadruple *tga2tga3tga5tga6* mutant displayed wild-type levels of cell death upon EE treatment, again suggesting that oxylipins do not play a role during this response. These results provide critical indications that HR-like triggered by *P. brassicae* eggs is independent from MYB30 and from oxylipin-mediated signaling pathways.

### Expression of sphingolipid metabolism genes

Sphingolipids are composed of a sphingoid LCB (long-chain base) backbone, produced by the condensation of a serine with a FA. LCB is then amidified to another FA moiety by ceramide synthases. A range of modifications can occur on LCB backbones such as hydroxylation or desaturation. These simple sphingolipids are named ceramides (Cer) and the usual nomenclature is to characterize them by both their LCB core structure as di- or tri-LCB (e.g. d18:0 for a dihydroxylated LCB with no unsaturation, t18:1 for a trihydroxylated LCB with one unsaturation and so on) and the FA moiety (e.g. C16:0 or h16:0 for 2-hydroxylated FA).

To explore the potential role of sphingolipids during egg-induced cell death, we measured the expression of different Arabidopsis genes involved in sphingolipid metabolism and signaling by qPCR after treatment with EE for 24, 48 and 72 h. Additionally, we performed the same analysis in *B. nigra* after 72 h, a plant species that was shown to develop HR-like lesions (Fatouros et al., 2014; Fatouros et al., 2016; Griese et al., 2017; Griese et al., 2021). Since the induction of cell death in *B. nigra* plants treated with EE was variable, in line with the phenotypes observed after natural oviposition on wild *B. nigra* constituting the original seed stock (Fatouros et al., 2014), we classified the response into weak symptom (HR-) or severe cell death (HR+) (Supplemental Figure S3). Remarkably, *LCB2b* and the ceramide synthase *LOH2* were consistently induced after 3 days of EE treatment in both plant species (Figure 4, Supplemental Figure S2 and S3). Interestingly, LOH2 catalyzes the attachment of C16:0 FA on dihydroxy LCB (d18:X), whereas LOH1 and LOH3 have a broader substrate specificity and attach mainly VLCFA on trihydroxy LCB (t18:X) (Luttgeharm et al., 2016; Ternes et al., 2011; Luttgeharm et al., 2015). Induction of *LOH2* thus suggests an increased metabolic flux towards C16-Cer (Figure 4), a class of known inducers of cell death in plants (Berkey et al., 2012).

**Figure 4.**
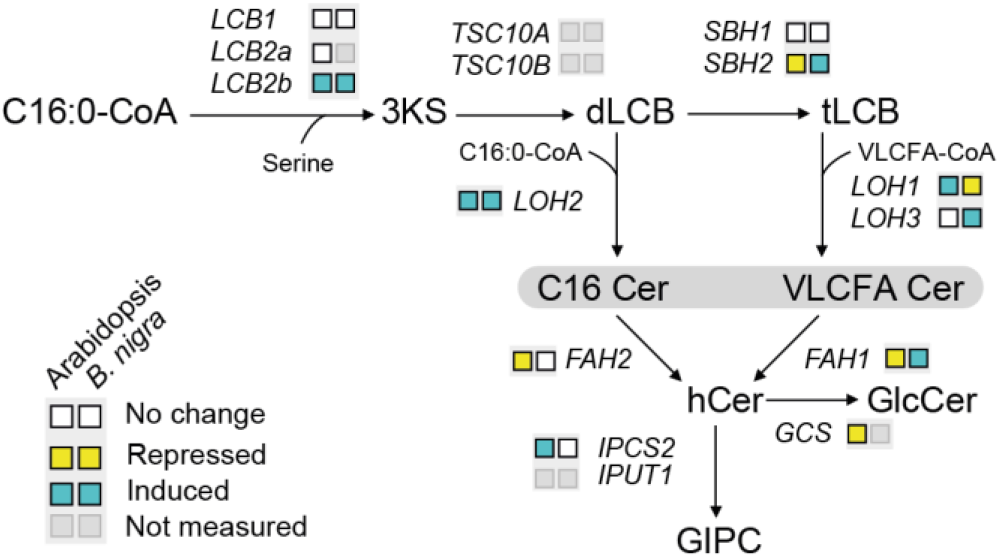
Expression of sphingolipid biosynthetic genes is altered upon EE treatment. Genes involved in each biosynthetic steps are indicated and substrates are indicated when appropriate. Results from expression analyses in Arabidopsis and *B. nigra* plants treated for three days with EE are indicated in color-coded boxes. Detailed expression data are available in Supplemental Figure S2 and S3. 3KS, 3-ketosphinganine; d/tLCB, di/tri-hydroxy long chain base; VLCFA, very-long chain fatty acid; Cer, ceramide; hCer, 2-hydroxyceramide; GlcCer, glucosylceramide; GIPC, glycosyl inositol phosphoryl ceramide.

Complex sphingolipids (GluCer and GIPC) consist for the main part of a trihydroxylated LCB attached to 2-hydroxy FA (hFA). The latter step is catalyzed by FAH1 and FAH2 (Nagano et al., 2012). Genes involved in fatty-acid hydroxylation (*FAH1/FAH2)* and GluCer synthesis (*GCS)* were downregulated in Arabidopsis (Supplemental Figure S2). This could indicate a decreased flux towards complex sphingolipids, possibly resulting in the accumulation of precursors (Cer and hCer). In contrast, *BnLOH3* and *BnFAH1* were induced in *B. nigra*, indicating a potential additional synthesis of complex sphingolipids in this species (Supplemental Figure S3). Altogether, these data further confirm that *P. brassicae* egg perception results in alterations of sphingolipid metabolism gene expression in two different plant species.

Although SA contributed partly to the expression of lipid metabolism genes (Figure 2C), *LOH2* and *LCB2b* were equally induced by EE in Col-0 and *sid2-1*, suggesting a SA-independent regulation of these genes (Supplemental Figure S2, B). Finally, to see whether the observed changes of sphingolipid metabolism gene expression after EE treatment might also occur during interaction with different types of attackers such as viruses, oomycetes, fungi, and bacteria, we explored publicly available transcriptome data from Genevestigator expression database (www.genevestigator.com). Interestingly, the pattern of sphingolipid-related gene expression was very similar between all biotic interactions, independently of the attacker or feeding mode considered (Supplemental Figure S4). This suggests that activation of sphingolipid metabolism gene expression is a conserved immune response.

### Ceramide synthase mutants show reduced EE-induced cell death

To further investigate the link between EE-triggered responses and sphingolipid metabolism, we tested whether cell death induction was altered in mutants lacking ceramide synthases LOH1, LOH2 or LOH3. Remarkably, both *loh2* and *loh3* displayed decreased cell death after three days of EE treatment, whereas *loh1* did not show any alteration (Figure 5A). These results are consistent with the observed induction of *LOH2* and supports a role for sphingolipids in the signaling pathway leading to EE-induced cell death.

**Figure 5.**
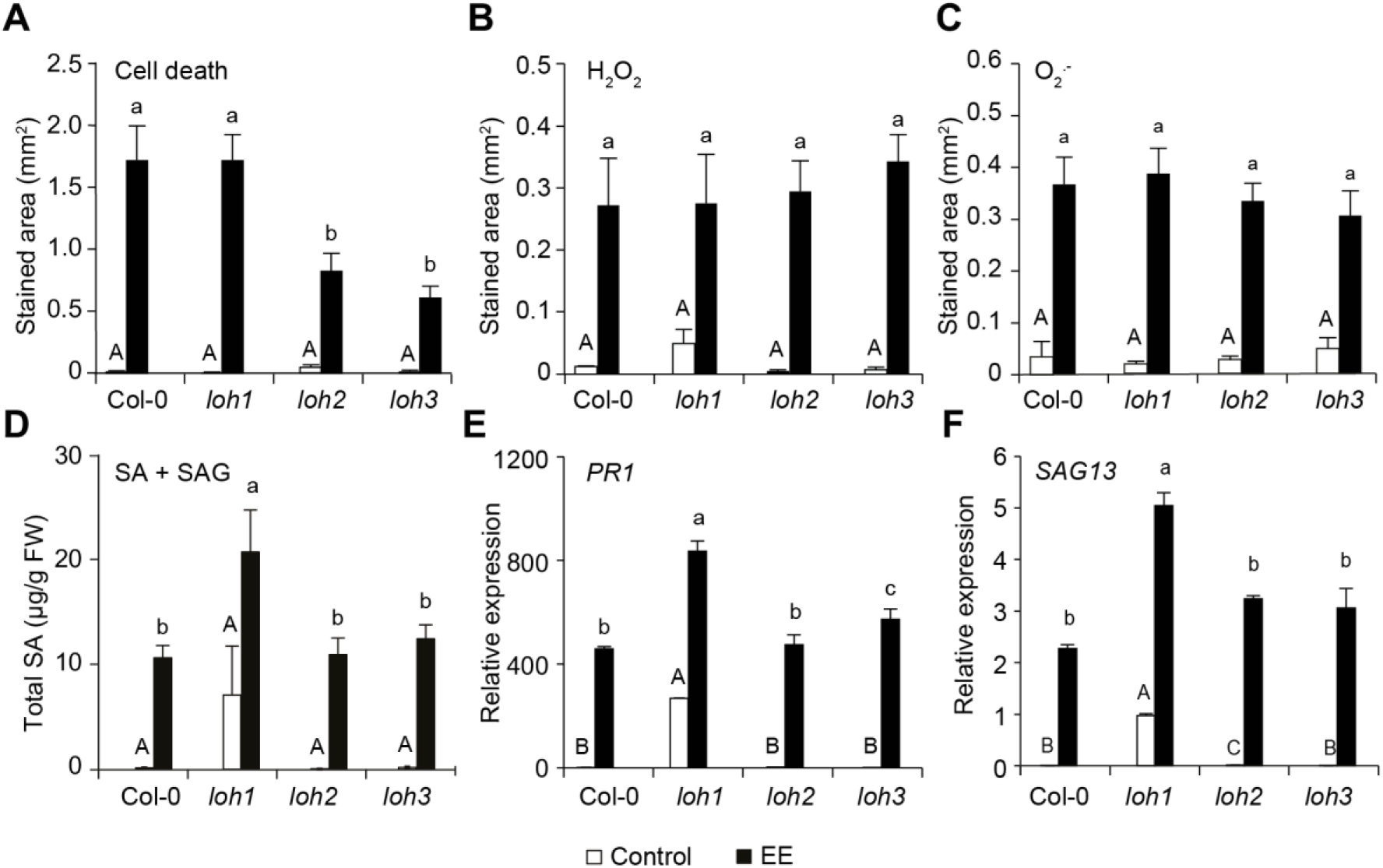
EE-induced cell death is reduced in ceramide synthase mutants. A-C, Cell death (A), H_2_O_2_ (B) or O ^.-^ (C) levels after three days of EE treatment in ceramide synthase mutants. For each genotype, a total of 12 to 16 leaves from 6 to 8 plants were treated with 2 μL of EE. Untreated leaves were used as controls. D, Total SA (SA + SAG) levels in ceramide synthase mutants after 3 days of EE treatment. Results from two independent experiments are shown (n=8). E,F, Expression of the SA-dependent marker *PR1* or the partially SA-dependent marker *SAG13* in ceramide synthase mutants after three days of EE treatment was monitored by qPCR. Data represent means ± SE of three technical replicates. Gene expression was normalized to the reference gene *SAND*. Different letters indicate significant differences at *P*<0.05 (ANOVA, followed by Tukey’s HSD for multiple comparisons). All experiments (except D) were repeated at least twice with similar results.

While studies have highlighted the existence of a link between SA signaling and sphingolipids, it is still unclear whether sphingolipids act upstream or downstream of SA accumulation and signaling during biotic interactions (Sánchez-Rangel et al., 2015). We previously reported that ROS and SA accumulation act as signals in response to insect eggs (Gouhier-Darimont et al., 2013; Gouhier-Darimont et al., 2019). We thus tested whether the lack of ceramide synthases in *loh2* and *loh3* affected the production of these early signals. Remarkably, no difference in H_2_O_2_ and O^2-^ production could be detected between Col-0 and mutant lines after EE treatment (Figure 5B,C), suggesting that LOH2 and LOH3 act downstream of ROS production.

Furthermore, SA reached similar levels after EE treatment in Col-0 and *loh2* or *loh3*, while *loh1* displayed higher basal SA levels and enhanced induction by EE treatment (Figure 5D). Moreover, EE-induced expression of the SA-dependent marker gene *PR1* and the partially SA-dependent marker *SAG13* was similar between Col-0 and *loh2* and *loh3* plants (Figure 5E,F). Consistent with higher SA levels, *loh1* displayed higher basal and induced transcript levels for these marker genes. Altogether, these results demonstrate a role for ceramides in the induction of cell death downstream of ROS and SA signaling.

### FA hydroxylation modulates EE-induced responses

2-hydroxylation of FA in ceramides is known to be crucial for complex sphingolipid synthesis (Markham et al., 2011; Ternes et al., 2011), and a link between 2-hydroxylation of FA and cell death was demonstrated (Nagano et al., 2012). The current model for sphingolipid synthesis suggests that α-hydroxylation occurs at the ceramide level through the activity of two isoforms of Fatty Acid Hydroxylase, FAH1 and FAH2 (König et al., 2012; Nagano et al., 2012). More specifically, FAH1 was shown to specifically hydroxylate VLCFA whereas FAH2 preferentially uses C16:0 FA as substrates. Interestingly, hVLCFA but not h16:0 FAs accumulated upon H_2_O_2_ treatment, suggesting that hVLCFA play a role in the suppression of cell death (Townley et al., 2005; Nagano et al., 2012). Based on our results demonstrating a role for ceramides in the induction of cell death following egg perception, we tested the contribution of sphingolipid FA hydroxylation in this response. After three days of EE treatment, cell death was slightly increased in the *fah1* mutant, but similar in *fah2* (Figure 6A). Further experiments revealed that while EE-induced H_2_O_2_ production was not altered (Figure 6B), basal and induced transcript levels of *PR1* and *SAG13* were higher in *fah1* than in Col-0 (Figure 6C and D). Collectively, these data suggest that hydroxylation of VLCFA in ceramides during egg-induced HR-like negatively regulates cell death and defense gene expression.

**Figure 6.**
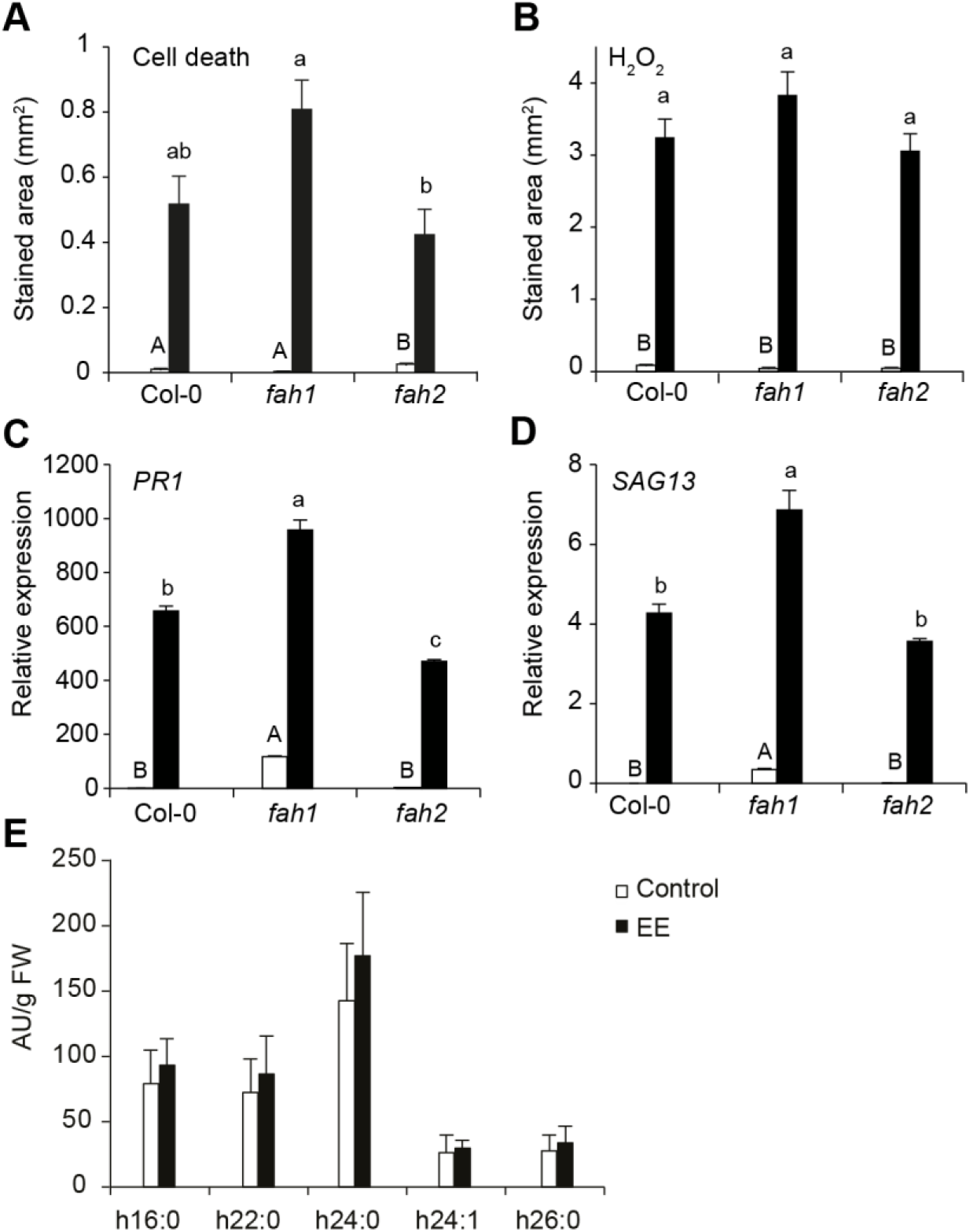
Fatty acid 2-hydroxylation modulates EE-induced cell death. A, Cell death and B, H_2_O_2_ levels after three days of EE treatments in fatty acid hydroxylase mutants. For each genotype, a total of 12 to 16 leaves from 6 to 8 plants were treated with 2 μL of EE. Untreated leaves were used as controls. C,D, Expression of the SA-dependent marker *PR1* and the partially SA-dependent marker *SAG13* after three days of EE treatment was monitored by qPCR. Data represent means ± SE of three technical replicates. Gene expression was normalized to the reference gene *SAND*. Different letters indicate significant differences at *P*<0.05 (ANOVA, followed by Tukey’s HSD for multiple comparisons). All experiments were repeated at least twice with similar results. E, 2-hydroxy fatty acid levels following three days of EE treatment were quantified by GC-MS. Data represent means ± SE from three biologically independent samples (n = 3).

We next examined FA 2-hydroxylation by GC-MS using a previously published method (Cacas et al., 2016). Surprisingly, we found that EE did not cause changes in the global distribution of hydroxy-FA levels (Figure 6E). One explanation could be that hydroxy-FA profiles from specific sphingolipid species may be altered while the overall amount remains stable.

### Sphingolipidome is altered in response to EE in Arabidopsis and *B. nigra*

Our results point to a role for C16:0-containing ceramides and hVLCFA-containing sphingolipids in modulating cell death in response to *P. brassicae* egg perception. However, the large number of sphingolipid species present in plants (> 200, Pata et al., 2010) renders the interpretation of phenotypes in sphingolipid-related mutants difficult. To resolve this issue, we determined the sphingolipidome composition of Arabidopsis and *B. nigra* plants in response to EE treatment. For the analysis, we used an extended LC-MS/MS analytical method that covers all sphingolipid classes, with the exception of phosphorylated Cer (Mamode Cassim et al., 2021). Additionally, to explore the link between sphingolipid alterations and SA (Sánchez-Rangel et al., 2015), we included *sid2-1* in our analysis.

Initial data exploration was performed by using 1-dimensional self-organizing map (1D-SOM) clustering to compare lipid profiles between Arabidopsis and *B. nigra* plants treated with EE (Figure 7A). Among the different clusters, several of them showed a pattern corresponding to an accumulation of lipids in response to EE in both (cluster 3 and 8) or in either plant species (cluster 2 for *B. nigra* and cluster 9 for Arabidopsis). Notably, cluster 3 and 8 contained three Cer 16:0 species as well as LCB t18:1, which are known cell death inducers, together with other Cer and GIPC (Supplemental Table S1). In contrast, cluster 2 was dominated by hFA-containing Cer and GIPC, while cluster 9 contained mainly non-hydroxy GIPC (Supplemental Table S1). Notably, no GluCer species were present in clusters correlating with HR-like cell death.

**Figure 7.**
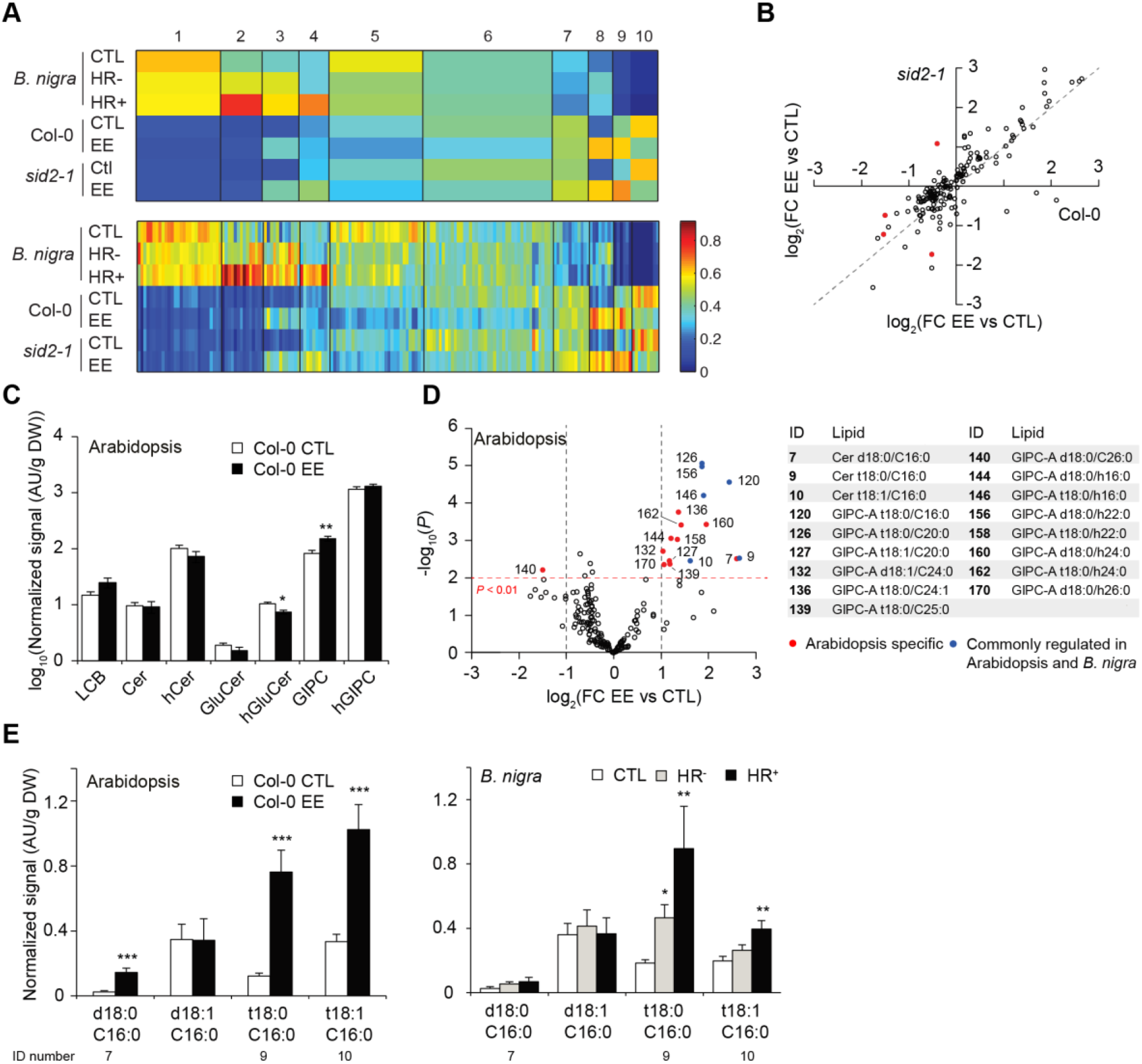
EE treatment induces changes in sphingolipid levels in both Arabidopsis and *B. nigra*. Leaves from either species were treated for three days with EE, and sphingolipids were extracted and analyzed by LC-MS/MS as described in methods. A, 1D-SOM clustering and heatmap visualizationof sphingolipid levels) using MarVis. Data were averaged over biological replicates (n=7), normalized using Euclidean unit length and the number of clusters was set to 10 (see Supplemental Table S4). The upper heatmap displays an average profile for each cluster and the one below displays all lipids individually. The list of markers found in each cluster can be found in Supplementable Table S4. B, Comparison of the impact of EE treatment on sphingolipid composition in Col-0 and *sid2-1* plants. Each circle represents one sphingolipid species. Filled circles are lipids whose level after EE treatment was significantly different between both genotypes; open circles are lipids whose level did not significantly change. The dotted line indicates perfect correspondence in accumulation between Col-0 and *sid2-1*. C, Levels of all major classes of sphingolipids in Col-0 plant treated or not with EE. Bars represent means ± SE from seven biologically independent samples (n = 7). D, Volcano plot of the sphingolipids detected in Col-0 plants. A threshold of P < 0.01 and a |FC| > 2 was used to identify molecules specifically changing upon EE treatment. Open circles indicate lipid species that did not significantly change, red and blue filled circles indicate lipid species that significantly changed upon EE treatment in *B.nigra* only or in both *B. nigra* and Arabidopsis respectively. ID for the latter is shown. A list of all significant lipids is shown on the right. E, Levels of the different C16-containing ceramides after EE treatment in Arabidopsis Col-0 plants (left) and *B. nigra* (right). Data represent means ± SE from four to ten independent samples. Asterisks denote statistically significant differences between EE treated samples and their respective controls (Welch *t*-test, *, *P*<0.05; **, *P*<0.01; ***, *P*<0.001). HR-, weak HR-like response; HR+, strong HR-like response.

Surprisingly, only a few lipids had a different accumulation pattern between Col-0 and *sid2-1* plants (Figure 7B), indicating that SA does not play a substantial role in sphingolipidome remodeling in response to EE. This conclusion is further supported by the fact that no cluster in Figure 7A shows a SA-dependent pattern. Globally, we could observe a significant increase and decrease in GIPC and hGluCer levels, respectively, following EE treatment in Arabidopsis (Supplemental Figure S10, S11), however no difference could be observed for the other classes (Figure 7C). In order to identify putative HR-like lipid markers, we next performed a volcano plot analysis on the sphingolipidome of Arabidopsis and *B. nigra*. Based on this analysis, we found 17 sphingolipids significantly accumulating in Col-0 plants following EE treatment (Figure 7D), while only one was less abundant. Top accumulating lipids in Arabidopsis were Cer C16:0 and both hydroxyl- and non-hydroxy GIPC (Supplemental Figure S7, S11). A number of markers were found to accumulate to a similar extent in *B. nigra,* including Cer t18:0/ and t18:1/C16:0 (Supplemental Figure S5). In addition, several GIPC, including GIPC t18:0/h16:0, also accumulated in both species. However, no obvious pattern regarding chain length or hydroxylation could be observed in both species (Supplemental Figure S8, S11, S12). Interestingly, while Arabidopsis significantly accumulated GIPC of all types (long chain and very long chain FA, hydroxylated or not) upon EE treatment, the response of *B. nigra* plants mainly showed an accumulation of GIPC C16:0 and h16:0 (Supplemental Figure S11A).

Overall, we observed common responses in both Arabidopsis and *B. nigra* following treatment with EE. Notably, we found an accumulation of several known cell death regulators, namely LCB t18:1 and Cer C16:0. In particular, the pattern of accumulation of Cer t18:0/ and t18:1/C16:0 was consistent with a role in cell death as shown by their strong accumulation in response to EE treatment in both species (Figure 7E). Moreover, in *B. nigra* this pattern correlated with HR intensity. In contrast, we could not observe such a pattern in LCB accumulation between both species, suggesting that they might not play a role in HR-like (Supplemental Fig S6). These results further support the hypothesis that EE treatment in Arabidopsis and *B. nigra* results in alterations of sphingolipid levels and a particular accumulation of Cer C16:0.

### Sphingolipidome of ceramide synthase mutants in response to EE

In order to investigate the role of individual ceramide synthases, we next quantified all sphingolipids in *loh1, loh2* and *loh3* mutant plants following EE treatment in a separate experiment. We observed that mutations in *loh1* and *loh2* had a larger impact on sphingolipidome remodeling after EE treatment as compared to *loh3* (Figure 8A-C). In particular, *loh1* plants displayed largely higher constitutive and induced levels of long chain FA-containing species from all sphingolipid classes (Supplemental Fig S12, S6-S11), making any comparison with other mutants difficult. In addition, this renders the identification of any EE-related cluster using 1D-SOM impossible when considering all mutants, since previously reported species constitutively over-accumulate in *loh1* (Supplemental Figure S12, Supplemental Table S5). Based on 1D-SOM clustering of the data from *loh2* and *loh3* only, we observed the existence of two clusters (cluster 1 and 2) that show an accumulation pattern consistent with a role in HR-like (Figure 8D). These clusters contained mostly C16:0 containing sphingolipids, including all Cer C16:0 and LCB t18:1 previously identified as potential HR-like regulators, further supporting our previous analysis. In particular, these clusters contained lipids present at very low levels in *loh2* plants, consistent with the previously reported catalytic activity of LOH2 and with the cell death phenotype observed. However, these lipids were still accumulating after EE treatment in *loh3* and no cluster could identify sphingolipids absent in both *loh2* and *loh3.* In line with this conclusion, we observed a significant accumulation of Cer d18:0/, t18:0/ and t18:1/C16:0 in Col-0 and *loh3* after upon treatment with EE which was absent in *loh2* mutant plants (Figure 8E). In contrast, *loh1* plants showed constitutive and induced levels of these lipids ∼10 fold higher than Col-0 plants and no further accumulation occurred after treatment. These results thus confirm the central role of LOH1 and LOH2 in sphingolipid metabolism and during the response to EE, but leaves the contribution of LOH3 unclear.

**Figure 8.**
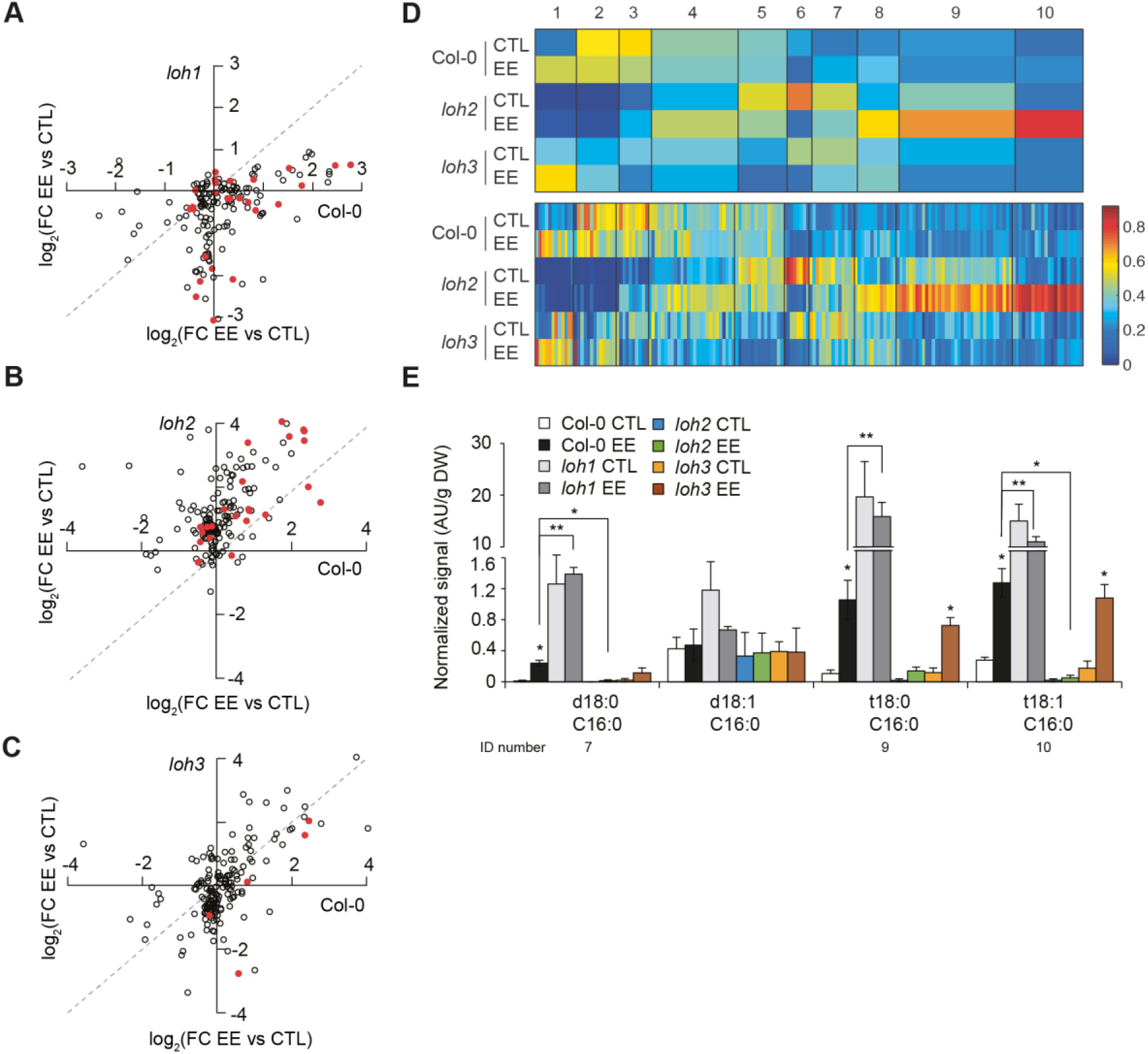
Sphingolipid levels in ceramide synthase mutants upon treatment with EE. Leaves were treated for three days with EE, and sphingolipids were extracted and analyzed by LC-MS/MS. A-C, Comparison of the impact of EE treatment on sphingolipid composition in Col-0 and *loh1* (A)*, loh2* (B) or *loh3* (C) mutant plants. Each circle represents one sphingolipid species. Filled circles are lipids whose level after EE treatment was significantly different between both genotypes; open circles are lipids whose level did not significantly change. The dotted line indicates perfect correspondence in accumulation between both genotypes. D, 1D-SOM clustering and heatmap visualization of sphingolipid levels using MarVis. Data were averaged over biological replicates (n=2-4), normalized using Euclidean unit length and the number of clusters was set to 10. The upper heatmap displays an average profile for each cluster and the one below displays all lipids individually. The list of lipids found in each cluster can be found in Supplemental Table S6. E, Levels of the different C16-containing ceramides after EE treatment in Arabidopsis Col-0 and *loh1 loh2* or *loh3* plants. Data represent means ± SE from two to four independent samples. Asterisks denote statistically significant differences between EE treated samples and their respective controls (Welch *t*-test, *, *P*<0.05; **, *P*<0.01; ***, *P*<0.001). Lines indicate statistically significant differences between Col-0 and mutant plants treated with EE.

## Discussion

We previously reported that the induction direct defenses against insect eggs in Arabidopsis involves the SA signaling pathway (Bruessow et al., 2010; Gouhier-Darimont et al., 2013). In addition, a growing body of evidence indicates that the response triggered by *P. brassicae* eggs are conserved in both *B. nigra* and Arabidopsis (Geiselhardt et al., 2013; Fatouros et al., 2014; Bonnet et al., 2017). However, components required for HR-like induction downstream of SA accumulation are still unknown. Besides their role as structural components of membranes, sphingolipids regulate PCD throughout the eukaryotic kingdom (Young et al., 2013). Analyzing previously published transcriptomic data on plant response to oviposition (Little et al., 2007), we observed an upregulation of genes related to sphingolipid metabolism. Experimental validation of this observation confirmed that EE triggers transcriptional alterations of genes involved in sphingolipid metabolism in both Arabidopsis and *B. nigra*. Furthermore, EE treatment led to the accumulation of various sphingolipid species. Despite differences in the profiles of other sphingolipids between Arabidopsis and *B. nigra*, a common and marked accumulation of Cer C16:0 and several GIPCs could be observed. Remarkably, the accumulation of Cer C16:0 is a conserved hallmark of cell death induction in plants, animals and fungi (Berkey et al., 2012; Young et al., 2013; Ali et al., 2018). In addition to the fact that EE was removed prior to sampling, most of the accumulating Cer C16:0 contained t18:0 or t18:1 LCB, which are not found in animals and GIPC are plant-specific lipids. It seems therefore unlikely that that the observed increase is due to the potential presence of egg-derived lipids. Interestingly, even though a similar accumulation of C16:0 was observed in response to *Botrytis cinerea* and *Pseudomonas syringae* pv. *tomato* AvrRPM1, the level of other sphingolipids, in particular GIPC, were largely different (Magnin-Robert et al., 2015).

We observed that cell death induction is dependent on the ceramide synthases LOH2 and LOH3, raising the question of the specificity of each enzyme during this process. Previous work revealed that LOH2 is responsible for the production of most C16:0-Cer *in planta*, while LOH1 and LOH3 were described as producing mainly VLCFA-Cer. Sphingolipid profiling in these mutants showed that *loh1* accumulated high levels of long chain-containing species whereas *loh2* plants were devoid of most of the C16:0 containing sphingolipids, consistent with previous studies (Ternes et al., 2011; Markham et al., 2011). *In vitro* ceramide synthase activity shows that LOH3, in contrast to LOH1, can also accept C16:0 substrates (Luttgeharm et al., 2016). We thus postulated that part of the observed accumulation of Cer C16:0 after EE treatment may originate from LOH3 activity in addition to LOH2. However, *loh3* mutants displayed mostly wild-type constitutive and induced levels of nearly all species detected, in agreement with previous studies (Ternes et al., 2011), thereby leaving the question of its contribution unclear. A contribution of VLCFA-containing sphingolipid to cell death was shown by the fact that *acd5loh2* double mutants have very low levels of C16:0-containing sphingolipids but still display spontaneous cell death (Bi et al., 2014). The accumulation of C16:0 in *loh1* mutants was previously linked to the appearance of spontaneous cell death after 8-10 weeks of development (Ternes et al., 2011). In our study, 4-5 weeks old *loh1* plants had high levels of Cer C16:0 yet did not display constitutive cell death, suggesting that plants trigger compensatory mechanisms to cope with the accumulation of these lipids before lesions appear. In addition, the context and the timing of this accumulation is largely different between development and biotic interactions, thereby complicating the interpretation of such data. These considerations together with the fact that only *loh2* plants showed a correlation between the levels of Cer C16:0 and the degree of cell death observed upon EE treatment may indicate that the potential role of Cer C16:0 during HR-like is complex. Alternatively, we cannot rule out that some of the observed phenotypes could be caused by non-catalytic activities of LOH and FAH genes. In agreement with this idea, FAH1 and FAH2 were previously shown to interact with the cell death suppressor BI-1 (Nagano et al., 2009).

How LCBs or ceramides induce cell death is still not clear (Berkey et al., 2012; De Bigault Du Granrut and Cacas, 2016) but studies from plants and other organisms may provide insights into their function. Sphingolipids, outside of their role as structural lipids in plasma membranes (PM), are major constituent of lipid nanodomains (Mamode Cassim et al., 2019). Membrane nanodomains, or so-called “lipid rafts”, are patches of lipids and proteins that laterally segregates from the rest of the PM due to high degree of intermolecular interactions between sphingolipids and sterols (Cacas et al., 2012; De Bigault Du Granrut and Cacas, 2016). Proteomic studies of nanodomain-associated proteins show that many immune regulators accumulate in these structures (Morel et al., 2006) and studies in rice showed that alterations in nanodomain sphingolipid composition can affect the abundance and function of PTI and cell death regulators in nanodomains (Ishikawa et al., 2015; Nagano et al., 2016). Additionally, phytoceramides were shown to perturb nanodomains in yeast (Pacheco et al., 2013). These results depict a dynamic relationship between sphingolipid metabolism and membrane nanodomains, suggesting that changes in the composition of certain sphingolipid classes could affect protein distribution and therefore nanodomain function.

In animal and yeast cells, ceramides have also been reported to have the ability to self-assemble so-called ceramide channels in mitochondria’s outer membrane and to promote the leakage of mitochondrial proteins (including cytochrome c), a hallmark point of no return for PCD (Siskind et al., 2002; Colombini, 2017). In this model, Cer are thus direct cell death executors, but whether such structures can form in plant cells is currently unknown. Interestingly, C16:0 phytoceramides (with trihydroxy LCBs) were reported to have a higher pore-forming activity in rat mitochondria than dihydroxy-ceramides (Perera et al., 2012), demonstrating that these molecules have the ability to form pores. Remarkably, these structures were found to assemble at physiological ceramide concentrations and their function was shown to be under the regulation of apoptotic regulators (Colombini, 2017). Alternatively, Cer may function through their interaction with cell-death modulator/executor proteins as revealed by the recent identification in human mitochondria of the voltage-dependent anion channel VDAC2 as a critical effector of Cer-induced PCD (Dadsena et al., 2019). In plants, no protein interacting with Cer have been described so far. However, the function of plant VDAC in PCD appears to be conserved when expressed in human cells, suggesting that Cer may also interact with VDAC channels in plants (Godbole et al., 2003). Given their important role in PCD induction in plants, future research should aim at identifying the molecular mechanisms underlying ceramide-regulated cell death.

Two recent studies in rice reported that alterations of hFA levels in nanodomains resulted in the depletion of certain PTI and cell death modulators (Ishikawa et al., 2015; Nagano et al., 2016). The role of sphingolipids in nanodomain structure and function may bring light on the observation that *fah1* displayed increased cell death following EE treatment, which correlated with a higher expression of SA-dependent markers *PR1* and *SAG13*. These data may thus suggest that the increased cell death observed in *fah1* after EE treatment may be the result of altered signaling from plasma membrane nanodomains. Although we found no clear alteration in overall hFA levels and distribution in different sphingolipid classes upon EE treatment, further in-depth work should examine the precise role of hFA species during HR-like.

Our results reveal a common accumulation of GIPC species in both Arabidopsis and *B. nigra* in response to EE perception. GIPC are critical regulators of membrane organization (Mamode Cassim et al., 2021). Studies show that GIPC accumulate in membrane nanodomains (Gronnier et al, 2016) and account for a large fraction of extracellular vesicles lipids (Liu et al., 2020), however the function of complex sphingolipids such as GlcCer and GIPC during biotic stresses is still poorly understood. Interestingly, it was recently shown that GIPC are receptors for fungal toxins (Lenarčič et al., 2017) and gate unknown Ca^2+^ channels upon binding of extracellular Na^+^ (Jiang et al., 2019). Whether GIPC play a role during HR-like is still not clear and work is needed to further explore this possibility.

The hypothesis that SA signaling and sphingolipid metabolism are somehow connected was based on the observation that certain sphingolipid mutants or treatment with the fungal toxin fumonisin B1 lead to increased defense gene expression or SA signaling/accumulation (Asai et al., 2006; Wang et al., 2008; König et al., 2012; Luttgeharm et al., 2015; Sánchez-Rangel et al., 2015; Fang et al., 2016). This led to postulate that the rise in Cer might precede SA accumulation (Sánchez-Rangel et al., 2015). We found that *loh2* and *loh3* mutant plants displayed wild-type ROS, SA and *PR1* levels, although they showed a significantly reduced cell death. These results thus suggest that ceramides function downstream of egg-induced SA signaling. However, we found no evidence that SA accumulation plays a role in the regulation of sphingolipid metabolism in the context of insect egg-triggered immunity as shown by the wild-type sphingolipid profile in *sid2-1* plants treated with EE. Additionally, we could not observe any significant alteration in the induction of *LOH2* and *LCB2b* in *sid2-1* compared to Col-0 after EE treatment, suggesting that SA does not contribute to the induction of these genes. It seems, however, that differences in sphingolipid-related gene expression may not necessarily always translate into changes in sphingolipid metabolism. This is evidenced by the differences observed in gene expression between Arabidopsis and *B. nigra* despite very similar sphingolipid accumulation patterns and by the fact that *B. nigra* HR-plants induced sphingolipid gene expression to a similar extent than HR+ plants, yet displaying intermediate Cer C16:0 accumulation. Thus, the regulation of sphingolipid accumulation in response to EE may be post-trancriptional.

Overall, our results show that sphingolipid metabolism plays a central role in the execution of HR-like in *Brassicaceae* after *Pieris brassicae* egg perception. Further research should investigate other plant-insect egg interactions and aim at deciphering the exact mechanism by which sphingolipids participate in HR-like.

## Materials and methods

### Plant and insect growth conditions

All experiments described using *Arabidopsis thaliana* were conducted in the Col-0 background. Seeds of *Brassica nigra* were collected from a wild population in Wageningen (The Netherlands) as previously described (Fatouros et al., 2014; Bonnet et al., 2017). Plants were grown in growth chambers in short day conditions (10 h light, 22°C, 65% relative humidity, 100 μmol m^-2^ s^-1^) and were 4 to 5 weeks old at the time of treatment. Seeds were stratified for 3 days at 4 °C after sowing. Larvae, eggs and butterflies of the Large White butterfly *Pieris brassicae* came from a population maintained on *Brassica oleracea* in a greenhouse as described previously (Reymond et al., 2000).

T-DNA insertion lines for *loh1* (SALK_069253), *loh2* (SALK_018608), *loh3* (SALK_150849), *fah1* (SALK_140660), *fah2* (SAIL_862_H01) were kindly provided by Ivo Feussner (University of Göttingen); *sid2-1* from Christiane Nawrath (University of Lausanne), *myb30* from Dominique Roby (LIPM INRA, Toulouse); *fad3fad7fad8* from Edward E. Farmer (University of Lausanne); *tga2tga3tga5tga6* from Corné Pieterse (Utrecht University).

### Whole-genome expression data

For *in silico* analysis of PCD (Figure 1) and lipid metabolism (Figure 2) marker gene expression, microarray data from Arabidopsis samples after 24 to 72 h of oviposition by *P. brassicae* (Little et al., 2007) were used. For *in silico* analysis of sphingolipid metabolism marker gene expression (Supplemental Figure S4), transcriptome data from Genevestigator public database (www.genevestigator.com) were used, except for data on *Pieris rapae* and *Spodoptera littoralis* feeding, which were from published microarray data (Reymond et al., 2004), and data on *P. brassicae* oviposition (Little et al., 2007).

### Treatment with egg extract

*P. brassicae* eggs were collected and crushed with a pestle in Eppendorf tubes. After centrifugation (15’ 000 g, 3 min), the supernatant (‘egg extract’, EE) was collected and stored at −20°C. Plants were 4-5 weeks old at the time of treatment. For each plant, two leaves were treated with 2 µl of EE. This amount corresponds to one egg batch of ca. 20 eggs. A total of four plants were used for each experiment. After the appropriate time, EE was gently removed with a scalpel blade and treated leaves were stored in liquid nitrogen. Untreated plants were used as controls.

### Histochemical staining

For visualization of cell death, EE was gently removed and leaves were submerged in lactophenol trypan blue solution (5 ml of lactic acid, 10 ml of 50% glycerol, 1 mg of trypan blue (Sigma), and 5 ml of phenol) at 28°C for 2–3 h. Hydrogen peroxide (H_2_O_2_) accumulation was measured with 3,3-diaminobenzidine (DAB; Sigma). Leaves were submerged in a 1.0 mg ml–1 DAB solution and incubated in the dark at room temperature for 6–8 h. Superoxide radical (O_2_.^−^) was visualized with the sensitive dye nitroblue tetrazolium (NBT; Sigma). Leaves were submerged in a solution containing 0.02% NBT and 10 mM NaN_3_ for 4 h at room temperature in the dark.

After each staining, leaves were destained for in boiling 95% ethanol. Microscope images were saved as TIFF files and processed for densitometric quantification with ImageJ version 1.64 (NIH).

### Salicylic acid quantification

SA quantifications were performed using the bacterial biosensor *Acinetobacter sp*. ADPWH (Huang et al., 2005; Huang et al., 2006) according to (DeFraia et al., 2008; Zvereva et al., 2016). Briefly, 6 leaf discs (0.7 cm, ∼20 mg) were ground in liquid nitrogen and extracted in 0.1M sodium acetate buffer (pH 5.6). Extracts were then centrifuged at 4°C for 15min at 16’000 g. 50 µL of extract were incubated with 5 µL of β-Glucosidase from almonds (0.5 U/µl in acetate buffer, Sigma-Aldrich) during 90 min at 37°C to release SA from SA-glucoside (SAG). 20 µL of extract was then mixed with 60 µL of LB and 50 µL of a culture of *Acinetobacter* sp. ADPWH_lux (OD_600_ = 0.4), and incubated for 1 h at 37°C. Finally, luminescence was integrated using a 485 ± 10 nm filter for 1 s. A 0 to 60 ng SA standard curve diluted in untreated *sid2-1* extract was read in parallel to allow quantification. SA amounts in samples were estimated by fitting a 3^rd^ order polynomial regression on the standards.

### Determination of non-enzymatic lipid peroxidation

Two leaves of each of three plants were treated for 3 days with EE and leaf discs (dia 0.7 cm) were harvested. Frozen leaf material (25 mg) was ground on liquid nitrogen, mixed with 0.5 ml of 0.1 % trichloroacetic acid (TCA), and centrifuged at 10’000 g for 15 min. 0.25 ml of the supernatant was mixed with 0.5 ml of 20 % TCA and 0.5 ml of 0.5 % thiobarbituric acid (TBA) and the mixture was incubated at 95 °C for 30 min to react MDA with TBA. Thereby a TBA-MDA complex is formed which is reported to have a specific absorbance at 532 nm. The specific absorbance of 532 nm and a nonspecific of 600 nm were measured with a UV-VIS spectrophotometer and the nonspecific absorbance was subtracted from the specific absorbance. The concentrations of MDA were calculated using Beer-Lambert’s equation with an extinction coefficient for MDA of 155 mM^-1^ cm^-1^ (Heath and Packer, 1968) and expressed to the fresh weight. Because this assay is described to overestimate the absolute concentration of MDA (Stahl et al., 2019), we normalized data on MDA levels in untreated Col-0 leaves and reported them as fold-changes.

### Hydroxy-fatty acid quantification

Hydroxy-FA quantification was performed based on a previously published protocol (Cacas et al., 2016). Briefly, 25 mg of frozen sample was spiked with 10 µg of heptadecanoic acid (C17:0) and 2-hydroxy-tetradecanoic acid (h14:0) as internal standards, and was transmethylated at 110°C overnight in 3 mL of methanol containing 5% (v/v) sulfuric acid. After cooling, 3 mL of NaCl (2.5%, w/v) was added, and methyl ester FAs were extracted in 1 mL of hexane. The organic phase was collected in a new tube, buffered with 3 mL of saline solution (200 mM NaCl and 200 mM Tris HCl, pH 8), centrifuged and the organic phase was dried under a gentle stream of nitrogen at room temperature. Free hydroxyl FAs were derivatized at 110°C for 30 min in 100 µL of N,O-bis(trimethylsilyl)trifluoroacetamide (BSTFA, Sigma) and pyridine (50:50, v/v), and surplus BSTFA was evaporated under nitrogen. Samples were finally dissolved in hexane and transferred into capped autosampling vials until analysis.

Quantitative analysis was performed using a HP-5MS capillary column (5% phenyl-methyl-siloxane, 30 m x 250 mm, 0.25 mm film thickness; Agilent) with helium carrier gas at 1.5 mL/min; injection was in splitless mode; injector temperature was set to 250°C; the oven temperature was held at 50°C for 1 min, then programmed with a 25°C/min ramp to 150°C (2 min hold), and a 10°C/min ramp to 320°C (6 min hold). Quantification of hydroxy-FAs was based on peak areas derived from specific ions in single-ion mode (SIM) and the respective internal standards. Ions used for quantifications are listed in the Supplemental Table S1.

### Sphingolipid analyses by LC-MS/MS

Analysis of sphingolipids by LC-MS/MS was done according to Mamode Cassim et al. (2021). Lipids extracts were incubated 1 h at 50°C in 2 mL of methylamine solution (7 ml methylamine 33% (w/v) in EtOH combined with 3 mL of methylamine 40% (w/v) in water (Sigma Aldrich) in order to hydrolyze phospholipids. After incubation, methylamine solutions dried at 40°C under a stream of air. Finally, were resuspended into 100 μL of THF/MeOH/H_2_O (40:20:40, v/v) with 0.1% formic acid containing synthetic internal lipid standards (LCB d17:1 [4.5 ng/μl], LCB-P d17:1 [4.5 ng/μl], Cer d18:1/C17:0 [4.5 ng/μl], GluCer d18:1/C12:0 [8.3 ng/μl] and GM1 [87 ng/μl], Avanti Polar Lipids) was added, thoroughly vortexed, incubated at 60°C for 20min, sonicated 2min and transferred into LC vials.

LC-MS/MS (multiple reaction monitoring mode) analyses were performed with a model QTRAP 6500 (ABSciex) mass spectrometer coupled to a liquid chromatography system (1290 Infinity II, Agilent). Analyses were performed in the positive mode. Nitrogen was used for the curtain gas (set to 30), gas 1 (set to 30), and gas 2 (set to 10). Needle voltage was at +5500 V with needle heating at 400°C; the declustering potential was adjusted between +10 and +40 V. The collision gas was also nitrogen; collision energy varied from +15 to +60 eV on a compound-dependent basis.

Reverse-phase separations were performed at 40°C on a Supercolsil ABZ+, 100 x 2.1 mm column and 5 µm particles (Supelco). The Eluent A was THF/ACN/5 mM Ammonium formate (3/2/5 v/v/v) with 0.1% formic acid and eluent B was THF/ACN/5 mM Ammonium formate (7/2/1 v/v/v) with 0.1% formic acid. The gradient elution program for LCB, Cer and GluCer quantification was as follows: 0 to 1 min, 1% eluent B; 40 min, 80% eluent B; and 40 to 42 min, 80% eluent B. The gradient elution program for GIPC quantification was as follows: 0 to 1 min, 15% eluent B;31 min, 45% eluent B; 47.5 min, 70% eluent B; and 47.5 to 49, 70% eluent B. The flow rate was set at 0.2 mL/min, and 5mL sample volumes were injected. A list of transitions for all sphingolipid species scanned during this procedure is available in Supplemental Table S2. The number of analyzed molecules per subclass scanned were: 4 LCB; 4 LCB-P; 110 Cer; 121 GluCer; 383 (64 GIPC serie A) GIPC.

The areas of LC peaks were determined using MultiQuant software (version 3.0; ABSciex) for sphingolipids quantification. Due to the lack of authentic standards for phytoceramides and GIPCs, the most abundant species present in plant tissues, absolute quantification is impossible without strong assumptions and was therefore avoided.

### Analysis of sphingolipid data

Areas of LC peaks for specific transitions were normalized to the signal of the standard from the same class (Cer, hCer, GluCer or GIPC) and normalized to sample dry weight. In total, 173 sphingolipid species could be quantified in all Arabidopsis and *B. nigra* samples (Supplemental Table S7). Data clustering and heatmaps were generated using the 1-dimensional self-organizing map (1D-SOM) algorithm implemented in the MarVis Cluster software (Kaever et al., 2009). Prior to clustering, replicate values were averaged and subsequent profiles were normalized to Euclidean unit length to allow comparison for all samples. The number of clusters was set to 10.

For volcano plots, comparisons between CTL and EE treated samples were performed using two-sided Welch T-test. For Arabidopsis datasets, initial data analysis was performed using a two-way ANOVA specifying genotype and treatment. *B. nigra* dataset was initially analyzed using a one-way ANOVA. Upon significant ANOVA analysis at *P* < 0.05, multiple comparisons were performed on the most informative sample pairs using two-sided Welch T-test without correction for multiple testing.

### RNA extraction, reverse-transcription and quantitative real-time PCR

Tissue samples were ground in liquid nitrogen, and total RNA was extracted using ReliaPrep™ RNA Tissue Miniprep (Promega) according to the manufacturer’s instructions, including DnaseI treatment. Afterwards, cDNA was synthesized from 500 ng of total RNA using M-MLV reverse transcriptase (Invitrogen) and subsequently diluted eightfold with water. Quantitative real-time PCR reactions were performed using Brilliant III Fast SYBR-Green QPCR Master Mix on an QuantStudio 3 real-time PCR instrument (Life Technologies) with the following program: 95°C for 3 min, then 40 cycles of 10 s at 95°C, 20 s at 60°C.

Values were normalized to the housekeeping gene *SAND* (At2g28390). The expression level of a target gene (TG) was normalized to the reference gene (RG) and calculated as normalized relative quantity (NRQ) as follows: NRQ = E^Ct^_RG_ /E^Ct^_TG_. For each experiment, three biological replicates were analyzed. A list of all primers used in experiments can be found in Supplemental Table S3.

Transcript sequences for gene homologs in *B. nigra* were identified by BLAST using Arabidopsis CDS sequences on BrassicaDB (http://brassicadb.org/brad/index.php). Because of the lack of accessible genome sequence, designed primer sequences were then checked for specificity using Primer BLAST against the “Brassica” mRNA database. Primer pairs that had no unspecific binding in other *Brassica* were tested by PCR on gDNA and cDNA from *B. nigra* for size and specificity.

### Statistics

Data were analyzed using R software version 3.6 or GraphPad Prism 9.0.1.

### Accession numbers

Sequence data from this article can be found in TAIR (www.arabidopsis.org) and BrassicaDB (http://brassicadb.org/brad/index.php) under the following accession numbers: *SAND* (At2g28390); *MYB30* (At3g28910); *FATB* (At1g08510); *PR1* (At2g14610); *SAG13* (At2g29350); *LOH1* (At3g25540); *LOH2* (At3g19260); *LOH3* (At1g13580); *LCB1* (At4g36480); *LCB2a* (At5g23670); *LCB2b* (At3g48780); *FAH1* (At2g34770); *FAH2* (At4g20870); *IPCS2* (At2g37940); *GCS* (At2g19880); *SBH1* (At1g69640); *SBH2* (At1g14290); *BnSAND* (BniB003645); *BnPR2* (BniB029818); *BnLOH1* (BniB046986); *BnLOH2* (BniB021107); *BnLOH3* (BniB004139); *BnLCB1* (BniB021240); *BnLCB2b* (BniB033113); *BnFAH1* (BniB033988); *BnFAH2* (BniB037392); *BnIPCS2* (BniB016748); *BnSBH1* (BniB037186); and *BnSBH2* (BniB033050).

## Supplemental data

The following materials are available in the online version of this article.

### Supplemental figures

**Supplemental Figure 1.**
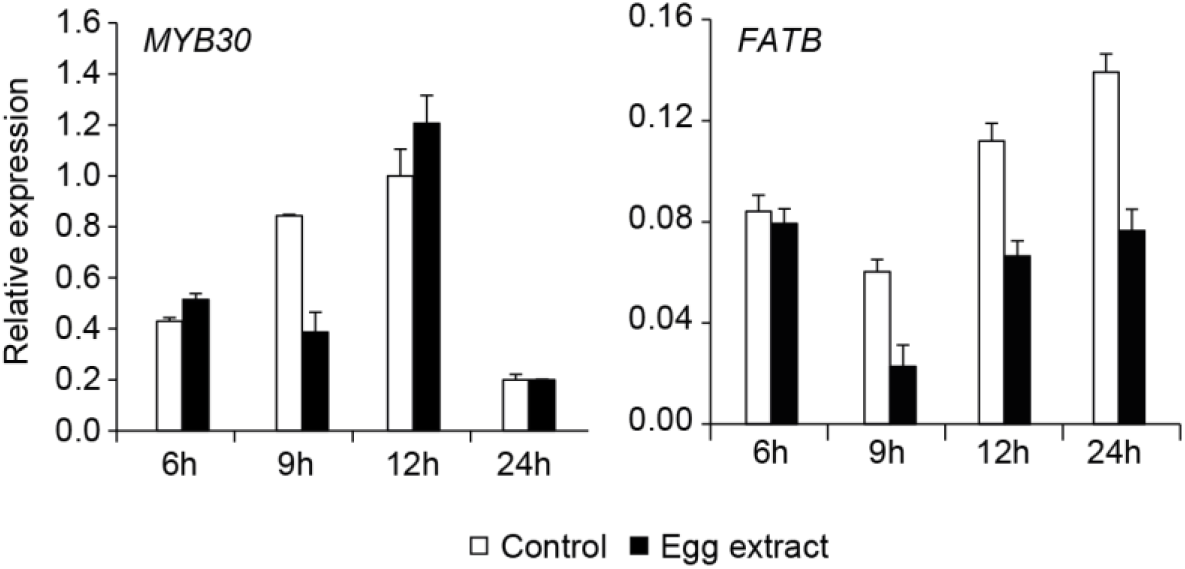
Expression of Arabidopsis *MYB30* and one of its target *FATB*. Expression after EE treatment was monitored by qPCR. Data represent means ± SE of three technical replicates. Gene expression was normalized to the reference gene *SAND*. This experiment was repeated once with similar results.

**Supplemental Figure 2.**
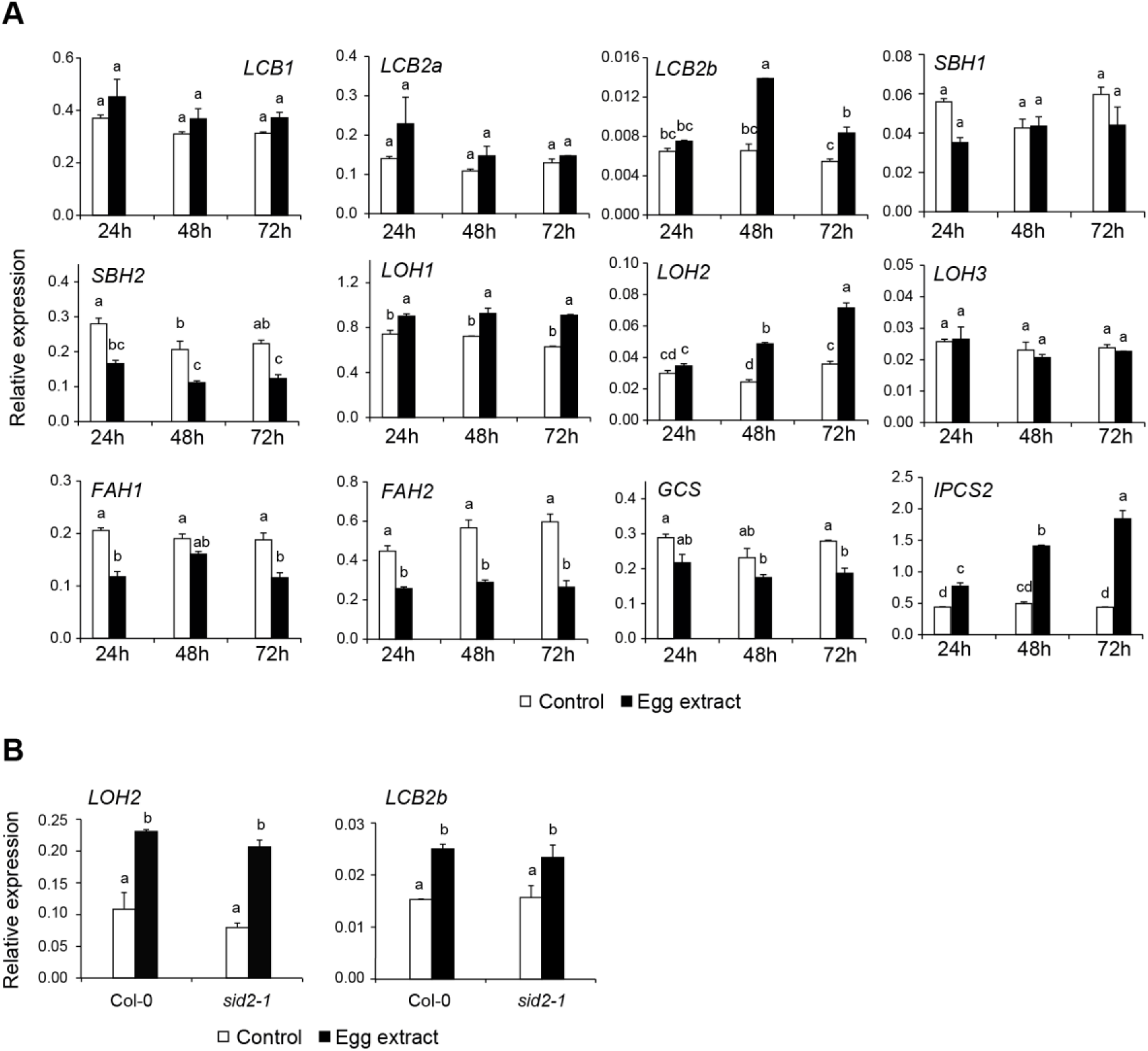
Time-course expression of sphingolipid metabolism genes in Arabidopsis. A, Expression of target genes was monitored by qPCR in Col-0 24 h to 72 h after EE treatment. B, Expression of target genes was monitored by qPCR in Col-0 and *sid2-1* 72 h after EE treatment. Data represent means ± SE of three technical replicates. Gene expression was normalized to the reference gene *SAND*. Different letters indicate significant differences at P<0.05 (ANOVA, followed by Tukey’s HSD for multiple comparisons). Experiments were repeated twice with similar results.

**Supplemental Figure 3.**
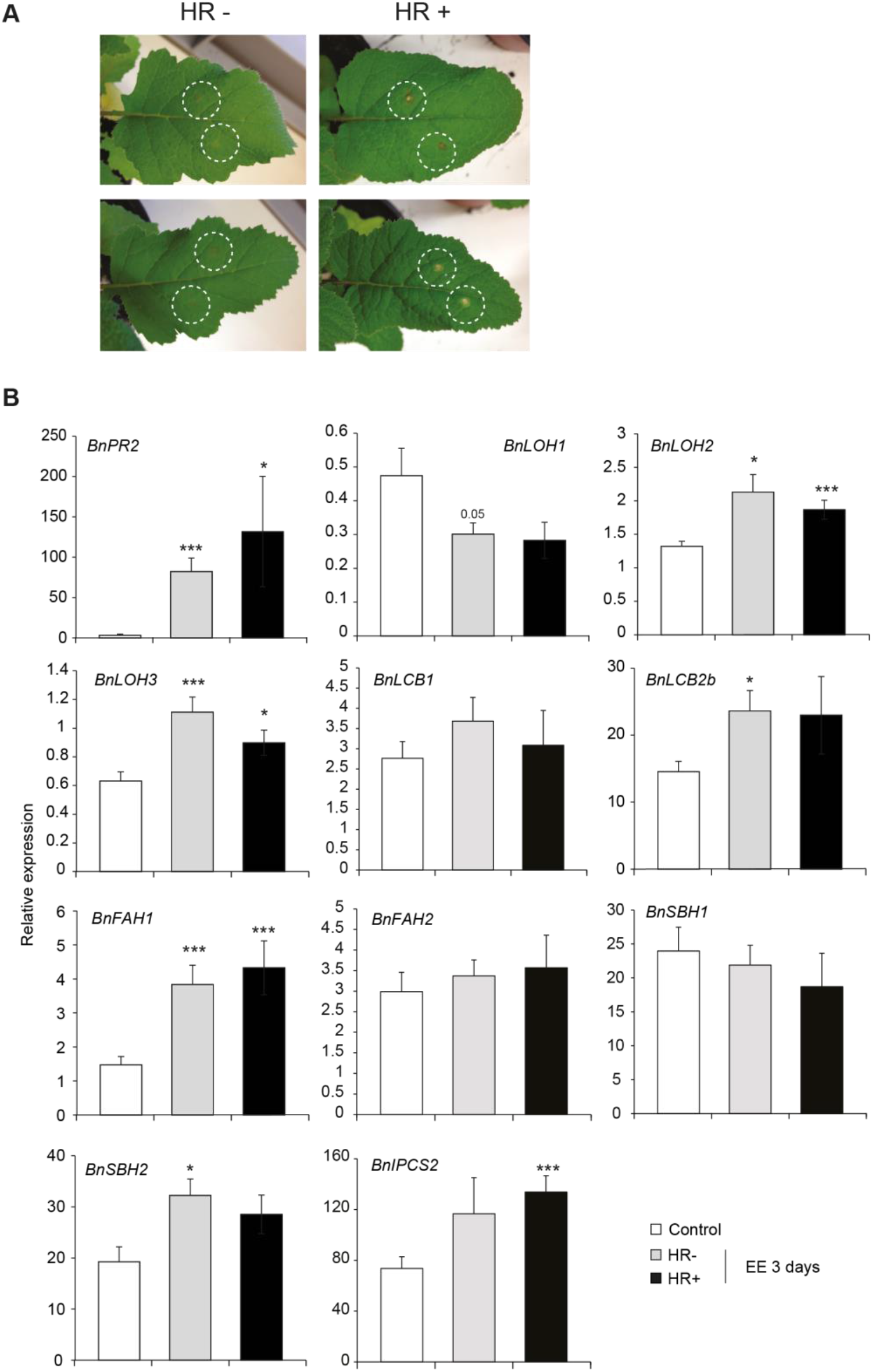
Time-course expression of sphingolipid metabolism genes in *Brassica nigr*a. A, Representative pictures of macroscopic HR-like symptoms triggered by *P. brassicae* EE after 3 days of treatment. Location of EE application is delineated by the white circles. B, Expression of target genes was monitored by qPCR 72 h after EE treatment. Gene expression was normalized to the reference gene *BnSAND*. Data represent means ± SE of three to seven independent biological replicates (n = 3-8). Asterisks denote statistically significant differences between EE treated and control plants (Welch *t*-test, *, P<0.05; **, P<0.01; ***, P<0.001). HR-, weak HR-like response; HR+, strong HR-like response.

**Supplemental Figure 4.**
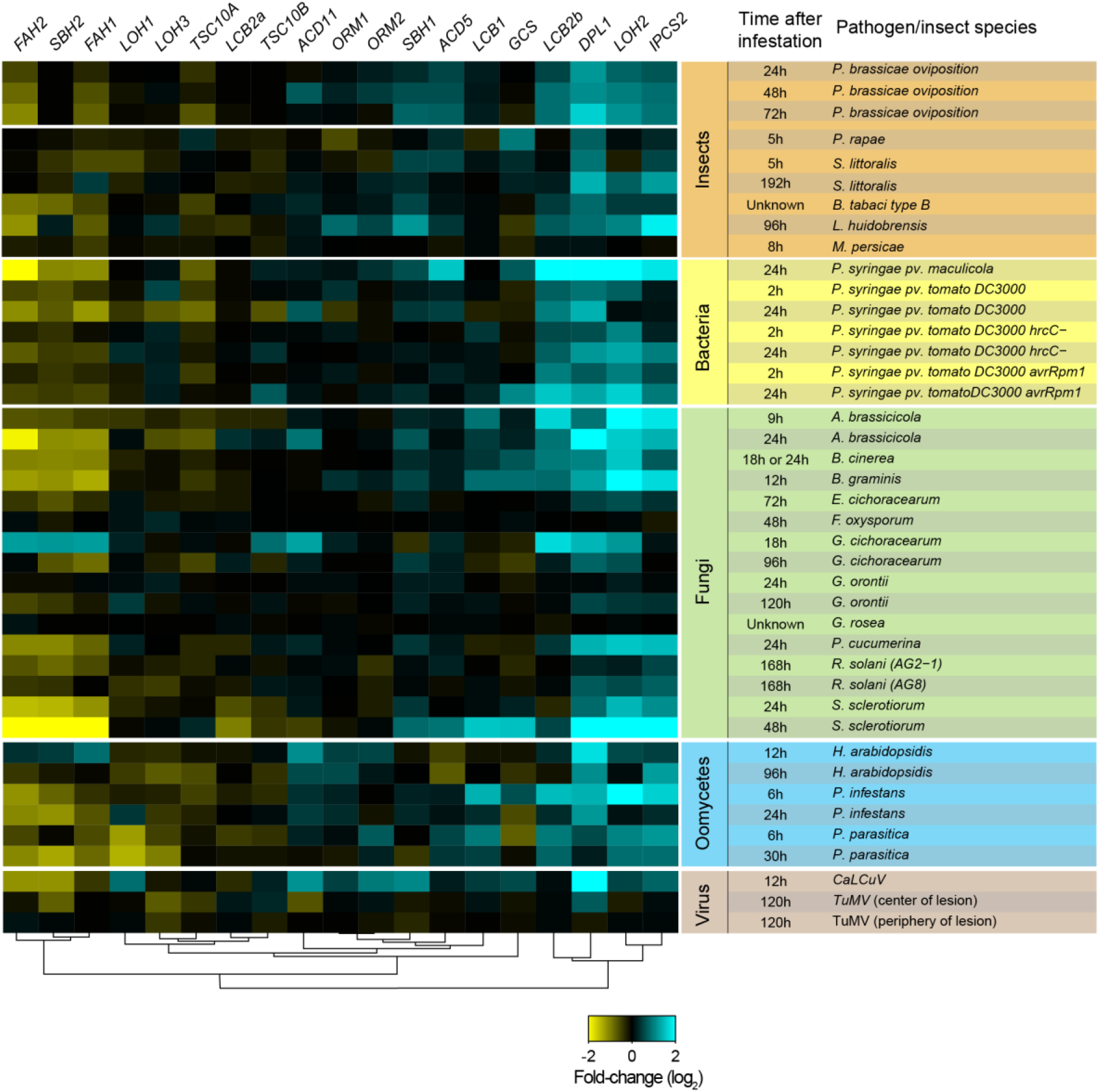
Diverse biotic stresses induce transcriptional alterations in sphingolipid metabolism. Relative expression levels of the selected sphingolipid genes in response to different attackers (insects, bacteria, fungi, oomycetes and virus) were obtained from Genevestigator. When more than two time-points were available, one early and one late time points were selected. Whole-genome expression data for *P. brassicae* oviposition or insect feeding (*Pieris rapae* or *Spodoptera littoralis* herbivory) were obtained from previous publications (see Methods for details).

**Supplemental Figure 5.**
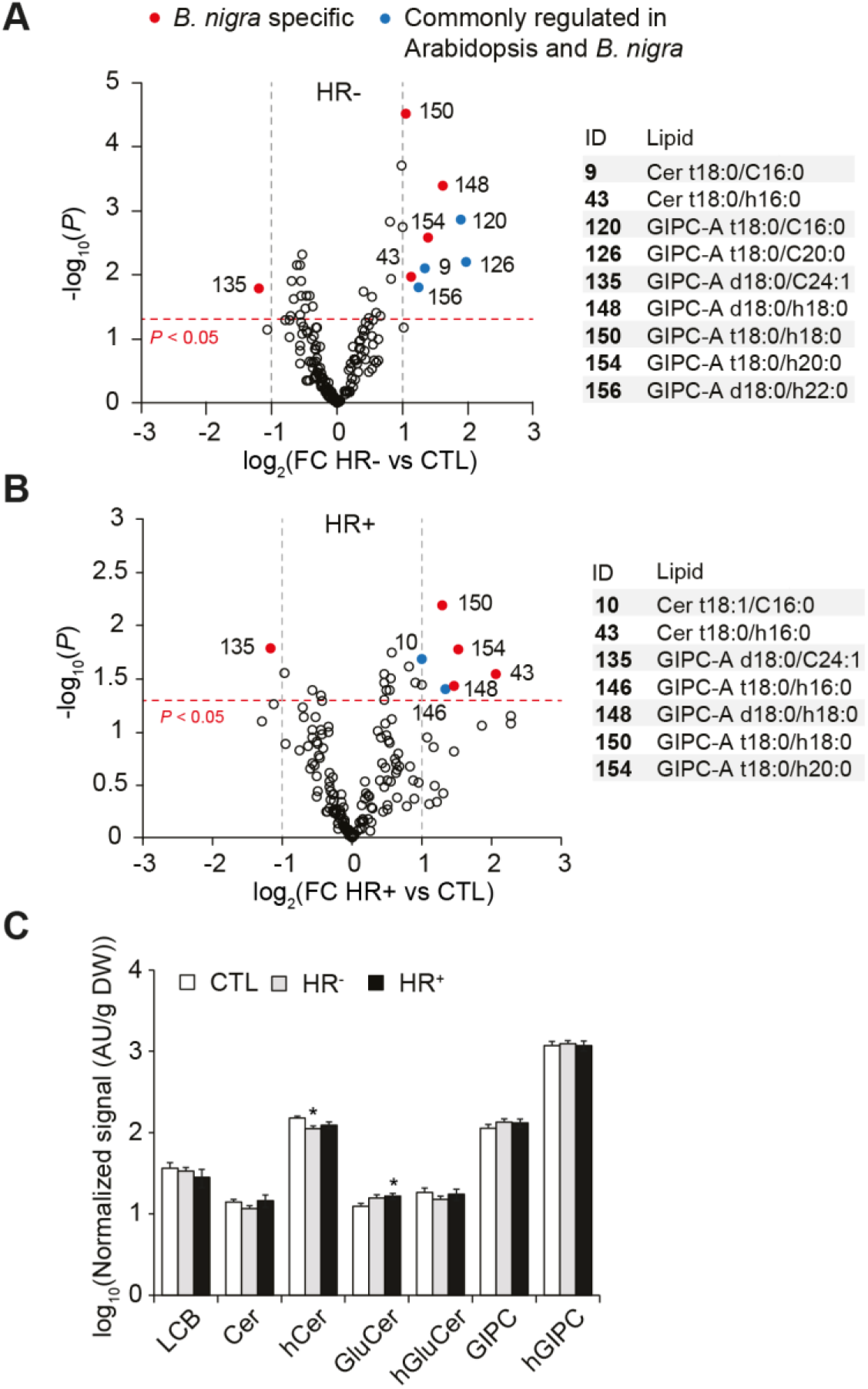
Effect of EE treatment in *B. nigra.* Leaves were treated for three days with EE, and sphingolipids were extracted and analyzed by LC-MS/MS. A,B, Volcano plot of the sphingolipids detected in HR- (A) and HR+ (B) *B. nigra* plants. A threshold of P < 0.05 and a |FC| > 2 was used to identify molecules specifically changing upon EE treatment. Open circles indicate lipid species that did not significantly change, red and blue filled circles indicate lipid species that significantly changed upon EE treatment in *B.nigra* only or in both *B. nigra* and Arabidopsis respectively. A list of all significant lipids is shown on the right. C, Levels of all major classes of sphingolipids in *B. nigra*. Bars represent means ± SE from seven biologically independent samples (n = 7). Data represent means ± SE from four to ten independent samples. Asterisks denote statistically significant differences between EE treated samples and their respective controls (Welch *t*-test, *, *P*<0.05; **, *P*<0.01; ***, *P*<0.001). HR-, weak HR-like response; HR+, strong HR-like response.

**Supplemental Figure 6.**
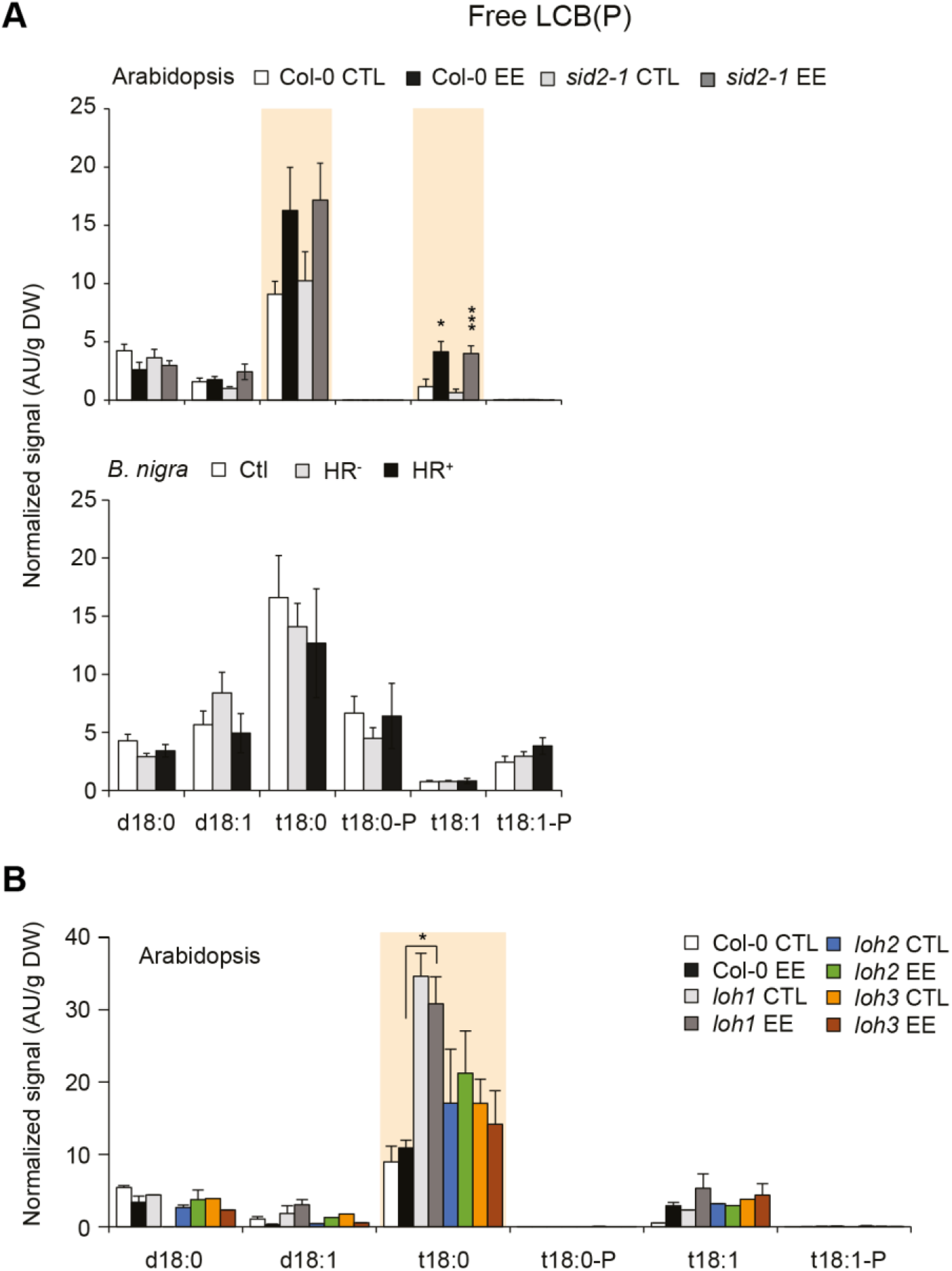
Free LCB and LCP-P levels in Arabidopsis wild-type and mutant lines and *B. nigra* plants following EE treatment. A, Col-0 and *sid2-1* mutant plants (upper panels) or *B. nigra* (lower panels) were treated with EE for three. Bars represent means ± SE from seven independent samples. B, Col-0, *loh1, loh2* and *loh3* mutant plants were treated as described in panel A. Bars represent means ± SE from two to four independent samples. Data were first analyzed using one-way (*B. nigra*) or two-way (Arabidopsis) ANOVA. Colored boxes indicate significant ANOVA at *P*< 0.05. Significant lipid markers were further analyzed using a selected number of pairwise comparisons: CTL and EE; EE-treated mutants compared to EE-treated Col-0. Asterisks denote statistical significance (Welch *t*-test between EE treated samples and their respective controls. *, *P*< 0.05; **, *P*< 0.01; ***, *P*<0.001). HR-, weak HR-like response; HR+, strong HR-like response.

**Supplemental Figure 7.**
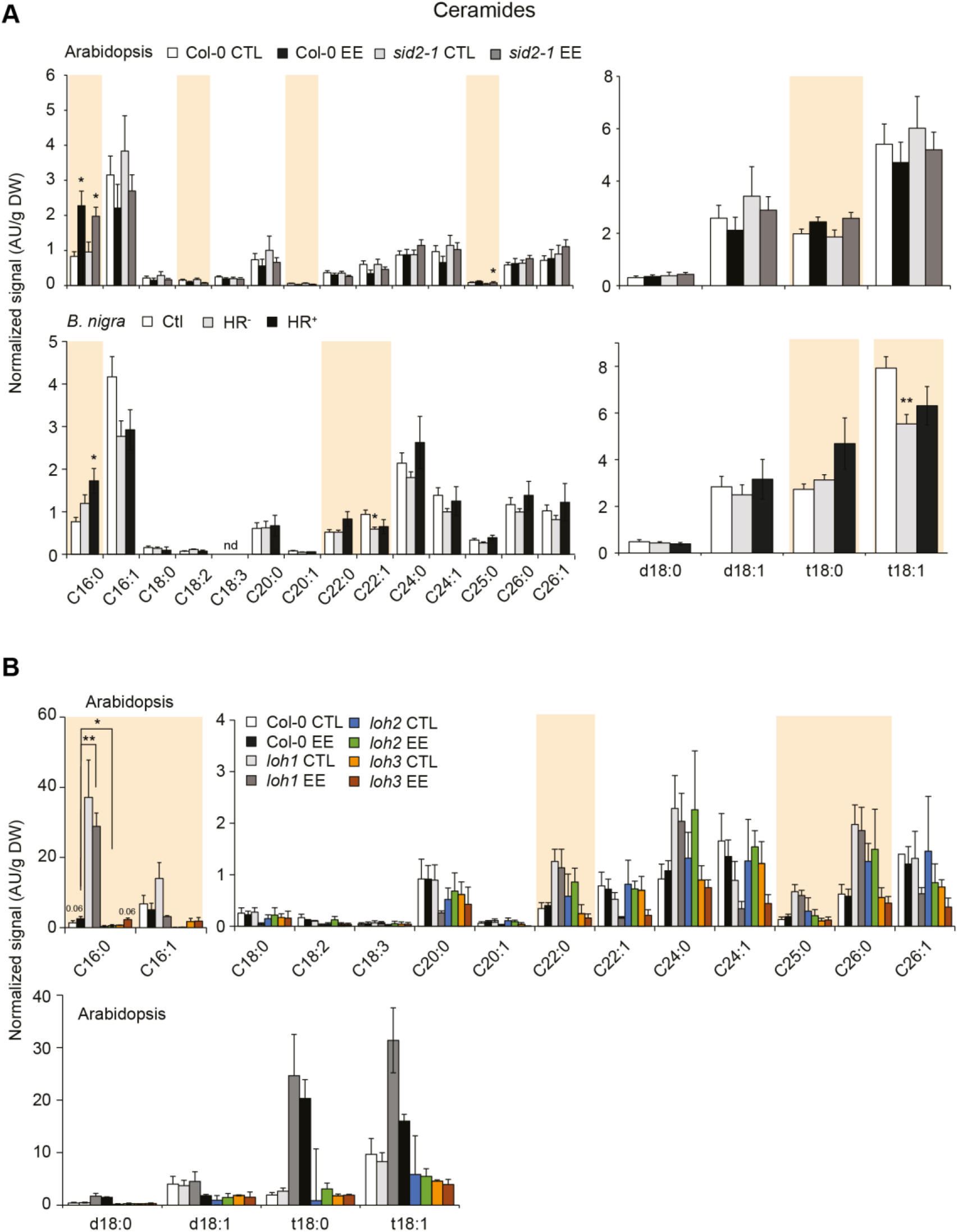
Ceramide levels in Arabidopsis wild-type and mutant lines and *B. nigra* plants following EE treatment. A, Col-0 and *sid2-1* mutant plants (upper panels) or *B. nigra* (lower panels) were treated with EE for three days. Sphingolipid levels are presented based on FA side-chain (left panel) or LCB (right panel) distribution. Bars represent means ± SE from seven independent samples. B, Col-0, *loh1, loh2* and *loh3* mutant plants were treated as described in panel A. Bars represent means ± SE from two to four independent samples. Data were first analyzed using one-way (*B. nigra*) or two-way (Arabidopsis) ANOVA. Colored boxes indicate significant ANOVA at *P* < 0.05. Significant lipid markers were further analyzed using a selected number of pairwise comparisons: CTL and EE; EE-treated mutants compared to EE-treated Col-0. Asterisks denote statistical significance (Welch *t*-test between EE treated samples and their respective controls. *, *P*< 0.05; **, *P*< 0.01). HR-, weak HR-like response; HR+, strong HR-like response.

**Supplemental Figure 8.**
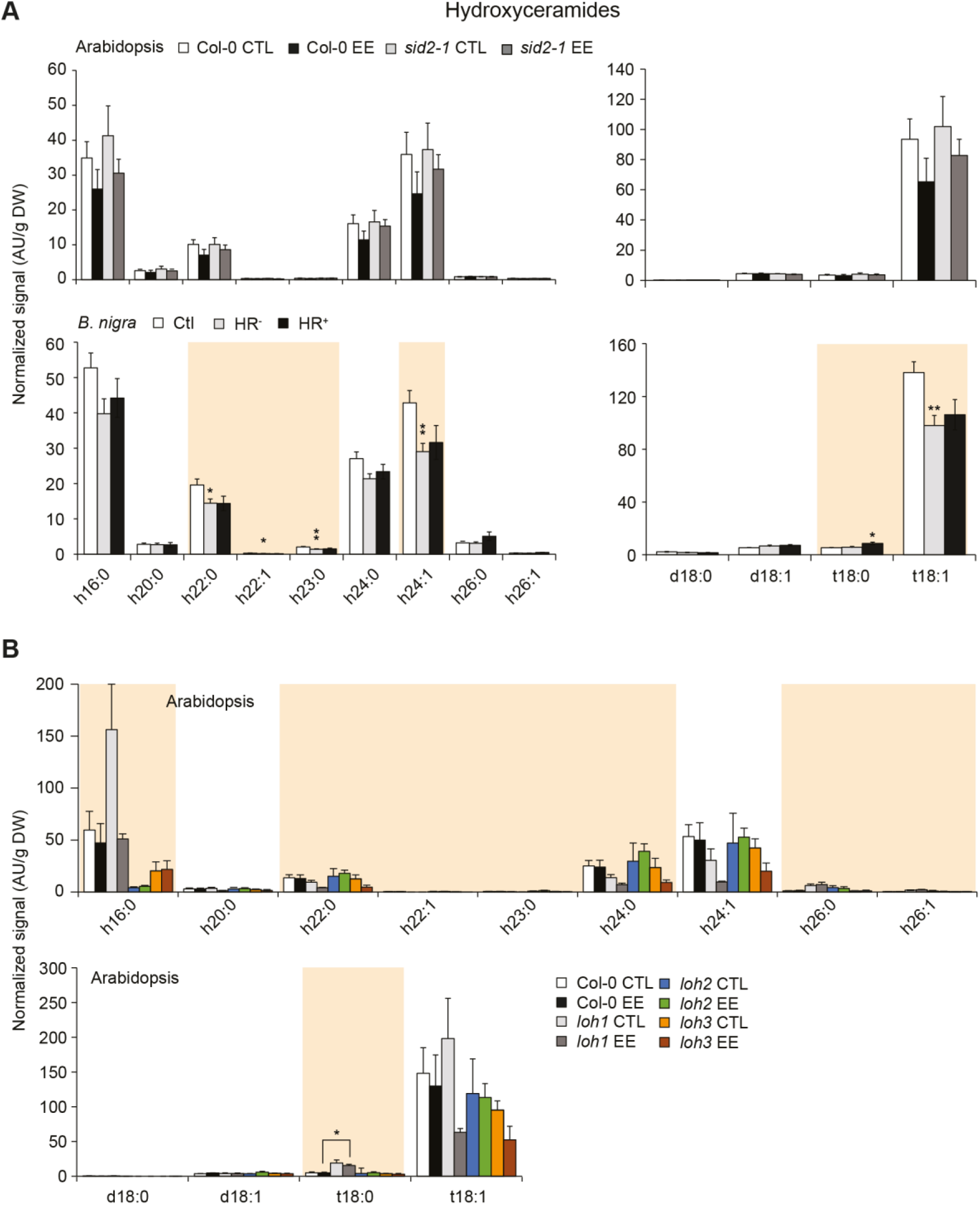
Hydroxy-ceramide levels in Arabidopsis wild-type and mutant lines and *B. nigra* plants following EE treatment. A, Col-0 and *sid2-1* mutant plants (upper panels) or *B. nigra* (lower panels) were treated with EE for three. Sphingolipid levels are presented based on FA side-chain (left panel) or LCB (right panel) distribution. Bars represent means ± SE from seven independent samples. B, Col-0, *loh1, loh2* and *loh3* mutant plants were treated as described in panel A. Bars represent means ± SE from two to four independent samples. Data were first analyzed using one-way (*B. nigra*) or two-way (Arabidopsis) ANOVA. Colored boxes indicate significant ANOVA at *P* < 0.05. Significant lipid markers were further analyzed using a selected number of pairwise comparisons: CTL and EE; EE-treated mutants compared to EE-treated Col-0. Asterisks denote statistical significance (Welch *t*-test between EE treated samples and their respective controls. *, *P*< 0.05; **, *P*< 0.01). HR-, weak HR-like response; HR+, strong HR-like response.

**Supplemental Figure 9.**
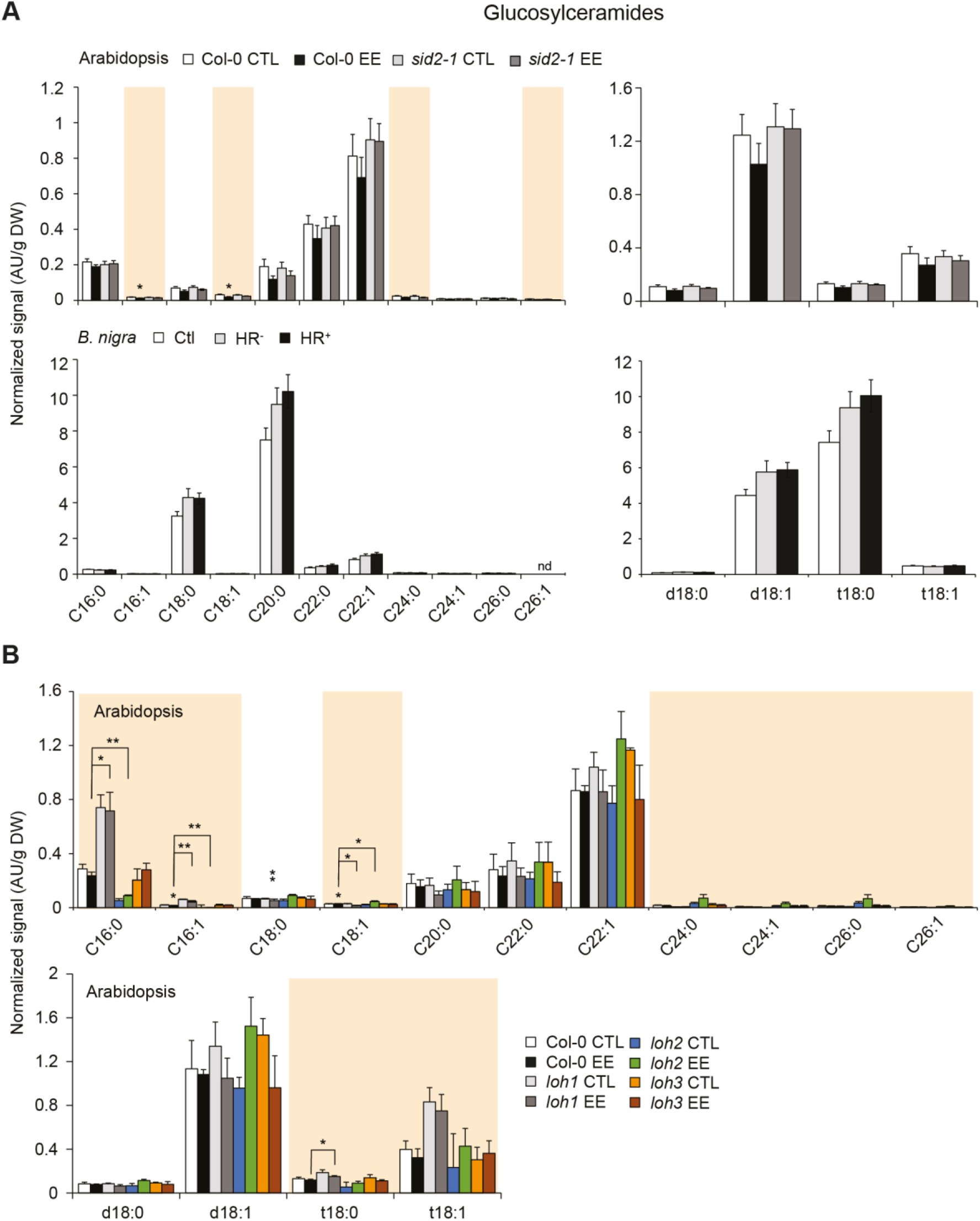
GluCer levels in Arabidopsis wild-type and mutant lines and *B. nigra* plants following EE treatment. A, Col-0 and *sid2-1* mutant plants (upper panels) or *B. nigra* (lower panels) were treated with EE for three days. Sphingolipid levels are presented based on FA side-chain (left panel) or LCB (right panel) distribution. Bars represent means ± SE from seven independent samples. B, Col-0, *loh1, loh2* and *loh3* mutant plants were treated as described in panel A. Bars represent means ± SE from two to four independent samples. Data were first analyzed using one-way (*B. nigra*) or two-way (Arabidopsis) ANOVA. Colored boxes indicate significant ANOVA at *P* < 0.05. Significant lipid markers were further analyzed using a selected number of pairwise comparisons: CTL and EE; EE-treated mutants compared to EE-treated Col-0. Asterisks denote statistical significance (Welch *t*-test between EE treated samples and their respective controls. *, *P*< 0.05; **, *P*< 0.01). HR-, weak HR-like response; HR+, strong HR-like response.

**Supplemental Figure 10.**
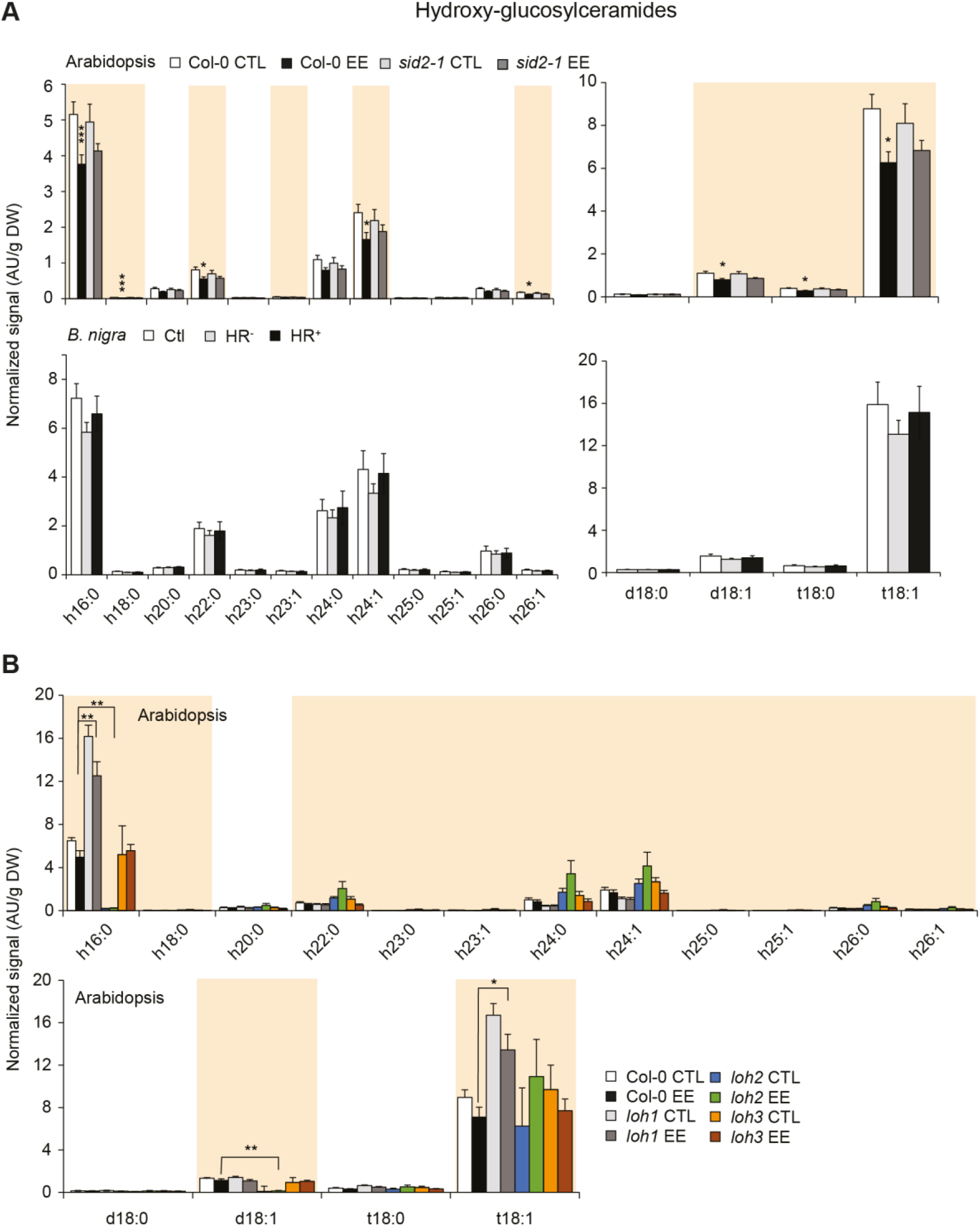
Hydroxy-GluCer levels in Arabidopsis wild-type and mutant lines and *B. nigra* plants following EE treatment. A, Col-0 and *sid2-1* mutant plants (upper panels) or *B. nigra* (lower panels) were treated with EE for three days. Sphingolipid levels are presented based on FA side-chain (left panel) or LCB (right panel) distribution. Bars represent means ± SE from seven independent samples. B, Col-0, *loh1, loh2* and *loh3* mutant plants were treated as described in panel A. Bars represent means ± SE from two to four independent samples. Data were first analyzed using one-way (*B. nigra*) or two-way (Arabidopsis) ANOVA. Colored boxes indicate significant ANOVA at *P* < 0.05. Significant lipid markers were further analyzed using a selected number of pairwise comparisons: CTL and EE; EE-treated mutants compared to EE-treated Col-0. Asterisks denote statistical significance (Welch *t*-test between EE treated samples and their respective controls. *, *P*< 0.05; **, *P*< 0.01; ***, *P*<0.001). HR-, weak HR-like response; HR+, strong HR-like response.

**Supplemental Figure 11.**
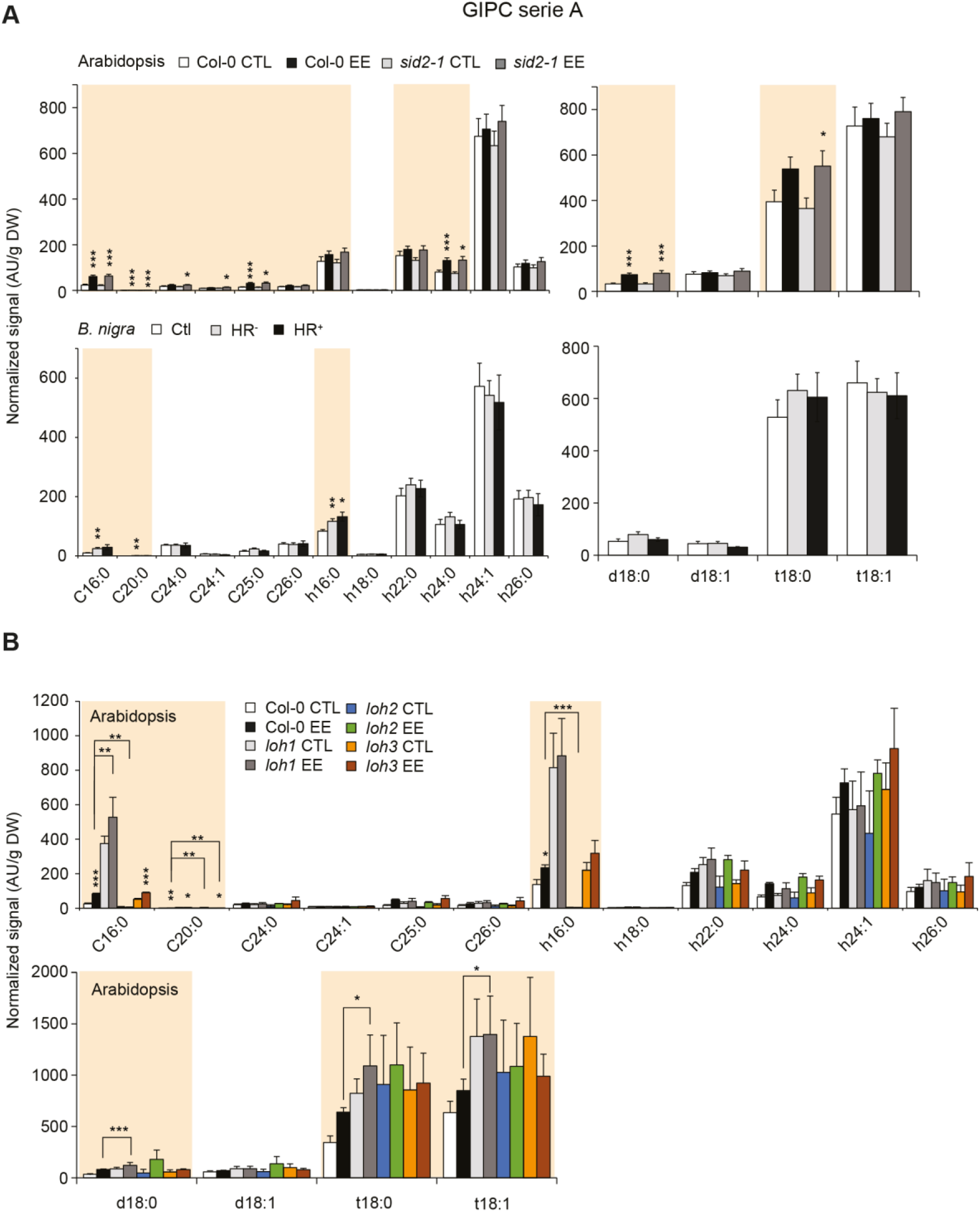
GIPC serie A levels in Arabidopsis wild-type and mutant lines and *B. nigra* plants following EE treatment. A, Col-0 and *sid2-1* mutant plants (upper panels) or *B. nigra* (lower panels) were treated with EE for three. Sphingolipid levels are presented based on FA side-chain (left panel) or LCB (right panel) distribution. Bars represent means ± SE from seven independent samples. B, Col-0, *loh1, loh2* and *loh3* mutant plants were treated as described in panel A. Bars represent means ± SE from two to four independent samples. Data were first analyzed using one-way (*B. nigra*) or two-way (Arabidopsis) ANOVA. Colored boxes indicate significant ANOVA at *P* < 0.05. Significant lipid markers were further analyzed using a selected number of pairwise comparisons: CTL and EE; EE-treated mutants compared to EE-treated Col-0. Asterisks denote statistical significance (Welch *t*-test between EE treated samples and their respective controls. *, *P*< 0.05; **, *P*< 0.01; ***, *P*<0.001). HR-, weak HR-like response; HR+, strong HR-like response.

**Supplemental Figure 12.**
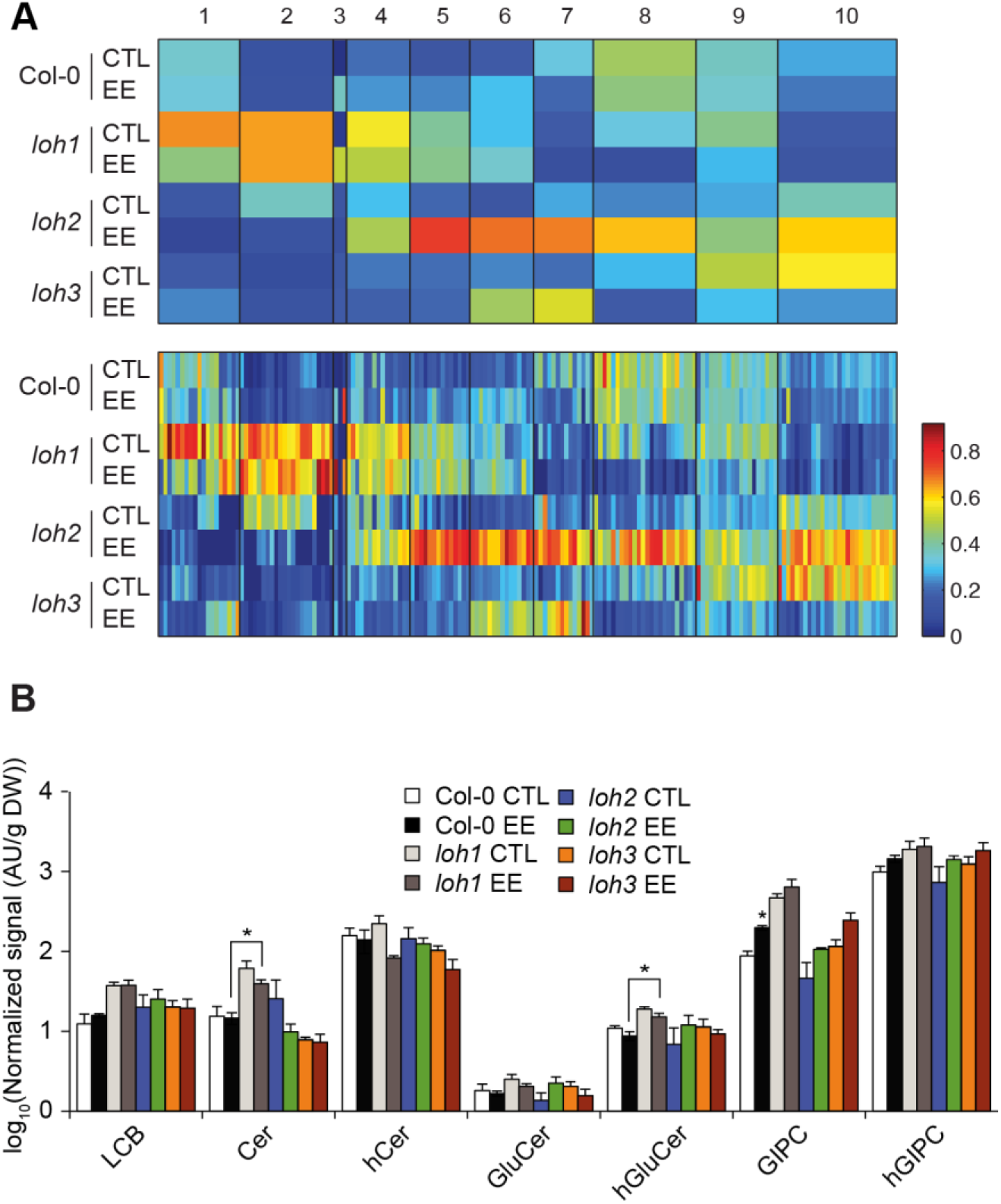
Effect of EE treatment on sphingolipid profiles in ceramide synthase mutants. Leaves from Col-0, *loh1, loh2* and *loh3* plants were treated for three days with EE. A, 1D-SOM clustering and heatmap visualization of sphingolipid levels) using MarVis. Data were averaged over biological replicates (n=2-4), normalized using Euclidean unit length and the number of cluster was set to 10. The upper heatmap displays an average profile for each cluster and the one below displays all lipids individually. The list of markers found in each cluster can be found in Supplemental Table S5. B, Levels of all major classes of sphingolipids. Bars represent means ± SE from two to four independent samples. Asterisks denote statistically significant differences between EE treated samples and their respective controls (Welch *t*-test, *, *P*<0.05).

**Supplemental Table S1.** List of ions used for FA and hFA identification and quantification by GC-MS.

**Supplemental Table S2.** List of all sphingolipid species targeted for LC-MS/MS analysis.

**Supplemental Table S3.** List of primers used for QPCR analyses.

**Supplemental Table S4.** 1D-SOM clustering results for Arabidopsis and *B. nigra* sphingolipid profiles.

**Supplemental Table S5.** 1D-SOM clustering results for Arabidopsis *loh1, loh2* and *loh3* sphingolipid profiles.

**Supplemental Table S6.** 1D-SOM clustering results for Arabidopsis *loh2* and *loh3* sphingolipid profiles.

**Supplemental Table S7.** Average individual sphingolipid levels from all experiments.

## Acknowledgments

We thank John Browse for sharing seeds of the *fad3 fad7 fad8* mutant. We also thank Blaise Tissot for maintenance of the plants; Steve Lassueur for pictures of HR-like symptoms on *B. nigra* leaves; Alice Bacon for valuable help with sphingolipid extractions; Marion Rebeaud, Lisa Butticaz and Dany Buffat for technical help during qPCR experiments. This work was supported by a grant from the Swiss National Science Foundation (grants no 31003A_169278 and 310030_200372 to P.R.). We thank Bordeaux-Metabolome platform for lipid analysis (www.biomemb.cnrs.fr/en/lipidomic-plateform/) supported by Bordeaux Metabolome Facility-MetaboHUB (grant no. ANR– 11–INBS–0010 to SM and LF).

## Author Contributions

R.G. and P.R. conceived the research plans. R.G. performed the experiments. L.F. and S.M. performed the sphingolipid analyses. R.G. and P.R. wrote the article with contributions of all authors. All authors read and approved the final version of the article.

## References

Ali U, Li H, Wang X, Guo L (2018) Emerging roles of sphingolipid signaling in plant response to biotic and abiotic stresses. Mol Plant 11: 1328–1343

Andersson MX, Hamberg M, Kourtchenko O, Brunnström Å, McPhail KL, Gerwick WH, Göbel C, Feussner I, Ellerström M (2006) Oxylipin profiling of the hypersensitive response in *Arabidopsis thaliana*: Formation of a novel oxo-phytodienoic acid-containing galactolipid, arabidopside E. J Biol Chem. 281: 31528– 31537

Asai T, Stone JM, Heard JE, Kovtun Y, Yorgey P, Sheen J, Ausubel FM (2000) Fumonisin B1-induced cell death in Arabidopsis protoplasts requires jasmonate-, ethylene-, and salicylate-dependent signaling pathways. Plant Cell 12: 1823–1836

Balbyshev N, Lorenzen J (1997) Hypersensitivity and egg drop: A novel mechanism of host plant resistance to Colorado potato beetle (Coleoptera: Chrysomelidae) J Econ Entomol 90: 652–657

Balint-Kurti P (2019) The plant hypersensitive response: Concepts, control and consequences. Mol Plant Pathol 20: 1163–1178

Begum MA, Shi XX, Tan Y, Zhou WW, Hannun Y, Obeid L, Mao C, Zhu Z-R (2016) Molecular characterization of rice *OsLCB2a1* gene and functional analysis of its role in insect resistance. Front Plant Sci. 7: 1789

Berkey R, Bendigeri D, Xiao S (2012) Sphingolipids and plant defense/disease: The “death” connection and beyond. Front Plant Sci 3: 68

Bonnet C, Lassueur S, Ponzio C, Gols R, Dicke M, Reymond P (2017) Combined biotic stresses trigger similar transcriptomic responses but contrasting resistance against a chewing herbivore in *Brassica nigra*. BMC Plant Biol 17: 127

Bruessow F, Gouhier-Darimont C, Bruchal A, Metraux, J-P, Reymond P (2010) Insect eggs suppress plant defence against chewing herbivores. Plant J 62: 876–885

Bruggeman Q, Raynaud C, Benhame M, Delarue, M (2015) To die or not to die? Lessons from lesion mimic mutants. Front Plant Sci 6: 1–22

Cacas JL, Furt F, Le Guédard M, Schmitter JM, Buré C, Gerbeau-Pissot P, Moreau P, Bessoule JJ, Simon-Plas F, Mongrand S (2012) Lipids of plant membrane rafts. Prog Lipid Res 51: 272–299

Cacas JL, Buré C, Grosjean K, Gerbeau-Pissot P, Lherminier J, Rombouts Y, Maes E, Bossard C, Gronnier J, Furt F, Fouillen L, et al. (2016) Revisiting plant plasma membrane lipids in tobacco: A focus on sphingolipids. Plant Physiol 170: 367–384

Carmona-Salazar L, Cahoon RE, Gasca-Pineda J, González-Solís A, Vera-Estrella R, Treviño V, Cahoon EB, Gavilanes-Ruiz M (2021) Plasma and vacuolar membrane sphingolipidomes: composition and insights on the role of main molecular species. Plant Physiol 186: 624–639

Coll NS, Epple P, Dangl JL (2011) Programmed cell death in the plant immune system. Cell Death Differ 18: 1247–56

Colombini M (2017) Ceramide channels and mitochondrial outer membrane permeability. J Bioenerg Biomembr 49: 57–64

Dadsena S, Bockelmann S, Mina JGM, Hassan DG, Korneev S, Razzera G, Jahn H, Niekamp P, Müller D, Schneider M, et al. (2019) Ceramides bind VDAC2 to trigger mitochondrial apoptosis. Nat Commun 10: 1832

De Bigault Du Granrut A, Cacas J-L (2016) How very-long-chain fatty acids could signal stressful conditions in plants? Front Plant Sci 7: 1–13

DeFraia CT, Schmelz EA, Mou Z (2008) A rapid biosensor-based method for quantification of free and glucose-conjugated salicylic acid. Plant Methods 4: 28

Fang L, Ishikawa T, Rennie EA, Murawska GM, Lao J, Yan J, Tsai AY, Baidoo EE, Xu J, Keasling JD (2016) Loss of inositol phosphorylceramide sphingolipid mannosylation induces plant immune responses and reduces cellulose content in Arabidopsis. Plant Cell 28: 2991–3004

Fatouros NE, Cusumano A, Danchin EGJ, Colazza S (2016) Prospects of herbivore egg-killing plant defenses for sustainable crop protection. Ecol Evol 6: 6906–6918

Fatouros NE, Pineda A, Huigens ME, Broekgaarden C, Shimwela MM, Candia IAF, Verbaarschot P, Bukovinszky T (2014) Synergistic effects of direct and indirect defences on herbivore egg survival in a wild crucifer. Proc R Soc B Biol Sci 281: 20141254

Fernandes GW (1990) Hypersensitivity: A neglected plant resistance mechanism against insect herbivores. Environ Entomol 19: 1173–1182

García-Marcos A, Pacheco R, Manzan, A, Aguilar E, Tenllado F (2013) Oxylipin biosynthesis genes positively regulate programmed cell death during compatible infections with the synergistic pair potato virus X-potato virus Y and tomato spotted wilt virus. J Virol 87: 5769–83

Garza R, Vera J, Cardona C, Barcenas N, Singh SP (2001) Hypersensitive response of beans to *Apion godmani* (Coleoptera: Curculionidae). J Econ Entomol 94: 958–962

Geiselhardt S, Yoneya K, Blenn B, Drechsler N, Gershenzon J, Kunze R, Hilker M (2013) Egg laying of Cabbage White butterfly (*Pieris brassicae*) on *Arabidopsis thaliana* affects subsequent performance of the larvae. PLoS One 8: e59661

Geuss D, Stelzer S, Lortzing T, Steppuhn A (2017) *Solanum dulcamara*’s response to eggs of an insect herbivore comprises ovicidal hydrogen peroxide production. Plant Cell Environ 40: 2663–2677

Godbole A, Varghese J, Sarin A, Mathew, M.K (2003) VDAC is a conserved element of death pathways in plant and animal systems. Biochim Biophys Acta - Mol Cell Res 1642: 87–96

Gouhier-Darimont C, Schmiesing A, Bonnet C, Lassueur S, Reymond P (2013) Signalling of *Arabidopsis thaliana* response to *Pieris brassicae* eggs shares similarities with PAMP-triggered immunity. J Exp Bot 64: 665–74

Gouhier-Darimont C, Stahl E, Glauser G, Reymond P (2019) The Arabidopsis lectin receptor kinase LecRK-I.8 is involved in insect egg perception. Front Plant Sci 10: 623

Griese E, Dicke M, Hilker M, Fatouros NE (2017) Plant response to butterfly eggs: inducibility, severity and success of egg-killing leaf necrosis depends on plant genotype and egg clustering. Sci Rep 7: 7316

Griese E, Caarls L, Bassetti N, Mohammadin S, Verbaarschot P, Bukovinszkine’Kiss G, Poelman EH, Gols R, Schranz ME, Fatouros NE (2021) Insect egg-killing: a new front on the evolutionary arms-race between brassicaceous plants and pierid butterflies. New Phytol 230: 341–353

Gronnier J, Germain V, Gouguet P, Cacas J-L, Mongrand S (2016) GIPC: glycosyl inositol phospho ceramides, the major sphingolipids on earth. Plant Signal Behav 11: e1152438

Heath RL, Packer L (1968) Photoperoxidation in isolated chloroplasts. I. Kinetics and stoichiometry of fatty acid peroxidation. Arch Biochem Biophys 125: 189–198

Hilfiker O, Groux R, Bruessow F, Kiefer K, Zeier J, Reymond P (2014) Insect eggs induce a systemic acquired resistance in Arabidopsis. Plant J 80: 1085–94

Huang WE, Huang L, Preston GM, Naylor M, Carr JP, Li Y, Singer AC, Whiteley AS, Wang H (2006) Quantitative in situ assay of salicylic acid in tobacco leaves using a genetically modified biosensor strain of *Acinetobacter* sp. ADP1. Plant J 46: 1073–1083

Huang WE, Wang H, Zheng H, Huang L, Singer AC, Thompson I, Whiteley AS (2005) Chromosomally located gene fusions constructed in *Acinetobacter* sp. ADP1 for the detection of salicylate. Environ Microbiol 7: 1339–1348

Huby E, Napier JA, Baillieul F, Michaelson LV, Dhondt-Cordelier S (2019) Sphingolipids: Towards an integrated view of metabolism during the plant stress response. New Phytol 225: 659–670

Huysmans M, Lema AS, Coll NS, Nowack MK (2017) Dying two deaths - programmed cell death regulation in development and disease. Curr Opin Plant Biol 35: 37–44

Ishikawa T, Aki T, Yanagisawa S, Uchimiya H, Kawai-Yamada M (2015) Overexpression of *BAX INHIBITOR-1* links plasma membrane microdomain proteins to stress. Plant Physiol 169: 1333–1343

Jiang Z, Zhou X, Tao M, Yuan F, Liu L, Wu F, Wu X, Xiang Y, Niu Y, Liu F, et al. (2019) Plant cell-surface GIPC sphingolipids sense salt to trigger Ca^2+^ influx. Nature 572: 341–346

Kaever A, Lingner T, Feussner K, Göbel C, Feussner I, Meinicke P (2009) MarVis: a tool for clustering and visualization of metabolic biomarkers. BMC Bioinformatics 10: 92

König S, Feussner K, Schwarz M, Kaever A, Iven T, Landesfeind M, Ternes P, Karlovsky P, Lipka V, Feussner I (2012) Arabidopsis mutants of sphingolipid fatty acid α-hydroxylases accumulate ceramides and salicylates. New Phytol 196: 1086– 1097

Lachaud C, Da Silva D, Amelot N, Béziat C, Brière C, Cotelle V, Graziana A, Grat S, Mazars C, Thuleau P (2011) Dihydrosphingosine-induced programmed cell death in tobacco BY-2 cells is independent of H_₂_O_₂_ production. Mol Plant 4: 310–8

Lenarčič T, Albert I, Böhm H, Hodnik V, Pirc K, Zavec AB, Podobnik M, Pahovnik D, Žagar E, Pruitt R, et al. (2017) Eudicot plant-specific sphingolipids determine host selectivity of microbial NLP cytolysins. Science 358: 1431–1434

Liang H, Yao N, Song JT, Luo S, Lu H, Greenberg JT (2003) Ceramides modulate programmed cell death in plants. Genes Dev 17: 2636–2641

Lim G, Singhal R, Kachroo A, Kachroo P (2017) Fatty acid– and lipid-mediated signaling in plant defense. Annu Rev Phytopathol 55: 505–536

Little D, Gouhier-Darimont C, Bruessow F, Reymond P (2007) Oviposition by pierid butterflies triggers defense responses in Arabidopsis. Plant Physiol 143: 784–800

Liu NJ, Wang N, Bao JJ, Zhu HX, Wang LJ and Chen XY (2020). Lipidomic analysis reveals the importance of GIPCs in Arabidopsis leaf extracellular vesicles. Mol Plant 13: 1523–1532.

Luttgeharm KD, Cahoon EB, Markham JE (2016) Substrate specificity, kinetic properties and inhibition by fumonisin B1 of ceramide synthase isoforms from Arabidopsis. Biochem J 473: 593–603

Luttgeharm KD, Chen M, Mehra A, Cahoon RE, Markham JE, Cahoon EB (2015) Overexpression of Arabidopsis ceramide synthases differentially affects growth, sphingolipid metabolism, programmed cell death, and mycotoxin resistance. Plant Physiol 169: 1108–17

Magnin-Robert, M, Le Bourse, D, Markham, J.E, Dorey, S, Clément, C, Baillieul, F, Dhondt-Cordelier, S (2015) Modifications of sphingolipid content affect tolerance to hemibiotrophic and necrotrophic pathogens by modulating plant defense responses in Arabidopsis. Plant Physiol 169: 2255–2274

Mamode Cassim A, Gouguet P, Gronnier J, Laurent N, Germain V, Grison M, Boutté Y, Gerbeau-Pissot P, Simon-Plas F, Mongrand S (2019) Plant lipids: Key players of plasma membrane organization and function. Prog Lipid Res 73: 1–27

Mamode Cassim A, Navon Y, Gao Y, Decossas M, Fouillen L, Grélard A, Nagano M, Lambert O, Bahammou D, Van Delft P, et al (2021) Biophysical analysis of the plant-specific GIPC sphingolipids reveals multiple modes of membrane regulation. J Biol Chem 296: 100602

Markham JE, Lynch DV, Napier JA, Dunn TM, Cahoon EB (2013) Plant sphingolipids: Function follows form. Curr Opin Plant Biol 16: 350–357

Markham JE, Molino D, Gissot L, Bellec Y, Hématy K, Marion J, Belcram K, Palauqui J-C, Satiat-Jeunemaître B, Faure J-D (2011) Sphingolipids containing very-long-chain fatty acids define a secretory pathway for specific polar plasma membrane protein targeting in Arabidopsis. Plant Cell 23: 2362–78

McConn M, Browse J (1996) The critical requirement for linolenic acid is pollen development, not photosynthesis, in an Arabidopsis mutant. Plant Cell 8: 403–416

Morel J, Claverol S, Mongrand S, Furt F, Fromentin J, Bessoule J-J, Blein J-P, Simon-Plas F (2006) Proteomics of plant detergent-resistant membranes. Mol Cell Proteomics 5: 1396–1411

Mueller S, Hilbert B, Dueckershoff K, Roitsch T, Krischke M, Mueller MJ, Berger S (2008) General detoxification and stress responses are mediated by oxidized lipids through TGA transcription factors in Arabidopsis. Plant Cell 20: 768–785

Nagano M, Ihara-Ohori Y, Imai H, Inada N, Fujimoto M, Tsutsumi N, Uchimiya H, Kawai-Yamada M (2009) Functional association of cell death suppressor, Arabidopsis Bax inhibitor-1, with fatty acid 2-hydroxylation through cytochrome b 5. Plant J 58: 122–134

Nagano M, Ishikawa T, Fujiwara M, Fukao Y, Kawano Y, Kawai-Yamada M, Shimamoto K (2016) Plasma membrane microdomains are essential for Rac1-RbohB/H-mediated immunity in rice. Plant Cell 28: 1966–1983

Nagano M, Takahara K, Fujimoto M, Tsutsumi N, Uchimiya H, Kawai-Yamada M (2012) Arabidopsis sphingolipid fatty acid 2-hydroxylases (AtFAH1 and AtFAH2) are functionally differentiated in fatty acid 2-hydroxylation and stress responses. Plant Physiol 159: 1138–1148

Olvera-Carrillo Y, Van Bel M, Van Hautegem T, Fendrych M, Huysmans M, Simaskova M, van Durme M, Buscaill P, Rivas S, Coll NS, et al. (2015) A conserved core of PCD indicator genes discriminates developmentally and environmentally induced programmed cell death in plants. Plant Physiol 169: 2684–2699

Pacheco A, Azevedo F, Rego A, Santos J, Chaves SR, Côrte-Real M, Sousa MJ (2013) C2-phytoceramide perturbs lipid rafts and cell integrity in *Saccharomyces cerevisiae* in a sterol-dependent manner. PLoS One 8: 1–12

Pata MO, Hannun YA, Ng CKY (2010) Plant sphingolipids: Decoding the enigma of the Sphinx. New Phytol 185: 611–630

Perera MN, Ganesan V, Siskind LJ, Szulc ZM, Bielawski J, Bielawska A, Bittman R, Colombini M (2012) Ceramide channels: Influence of molecular structure on channel formation in membranes. Biochim Biophys Acta - Biomembr 1818: 1291–1301

Petzold-Maxwell J, Wong S, Arellano C, Gould F (2011) Host plant direct defence against eggs of its specialist herbivore, *Heliothis subflexa*. Ecol Entomol 36: 700– 708

Radojičić A, Li X, Zhang Y (2018) Salicylic acid: A double-edged sword for programed cell death in plants. Front Plant Sci 9: 1133

Raffaele S, Vailleau F, Leger A, Joubes J, Miersch O, Huard C, Blee E, Mongrand S, Domergue F, Roby D (2008) A MYB transcription factor regulates very-long-chain fatty acid biosynthesis for activation of the hypersensitive cell death response in Arabidopsis. Plant Cell 20: 752–767

Reape TJ, McCabe PF (2010) Apoptotic-like regulation of programmed cell death in plants. Apoptosis 15: 249–256

Reymond P (2013) Perception, signaling and molecular basis of oviposition-mediated plant responses. Planta 238: 247–58

Reymond P, Bodenhausen N, Van Poecke RMP, Krishnamurthy V, Dicke M, Farmer EE (2004) A conserved transcript pattern in response to a specialist and a generalist herbivore. Plant Cell 16: 3132–3147

Salguero-Linares J, Coll NS (2019) Plant proteases in the control of the hypersensitive response. J Exp Bot 70: 2087–2095

Salvesen GS, Hempel A, Coll NS (2015) Protease signaling in animal and plant regulated cell death. FEBS J 283: 2577–2598

Sánchez-Rangel D, Rivas-San Vicente M, de la Torre-Hernández ME, Nájera-Martínez M, Plasencia J (2015) Deciphering the link between salicylic acid signaling and sphingolipid metabolism. Front Plant Sci 6: 1–8

Saucedo-García M, Guevara-García A, González-Solís A, Cruz-García F, Vázquez-Santana S, Markham JE, Lozano-Rosas MG, Dietrich CR, Ramos-Vega M, Cahoon EB, et al. (2011) MPK6, sphinganine and the *LCB2a* gene from serine palmitoyltransferase are required in the signaling pathway that mediates cell death induced by long chain bases in Arabidopsis. New Phytol 191: 943–957

Shapiro AM, DeVay JE (1987) Hypersensitivity reaction of *Brassica nigra* L. (Cruciferae) kills eggs of *Pieris* butterflies (Lepidoptera: Pieridae). Oecologia 71: 631–632

Shi L, Bielawski J, Mu J, Dong H, Teng C, Zhang J, Yang X, Tomishige N, Hanada K, Hannun YA, et al. (2007) Involvement of sphingoid bases in mediating reactive oxygen intermediate production and programmed cell death in Arabidopsis. Cell Res 17: 1030–1040

Siebers M, Brands M, Wewer V, Duan Y, Hölzl G, Dörmann P (2016) Lipids in plant– microbe interactions. Biochim Biophys Acta - Mol Cell Biol Lipids 1861: 1379– 1395

Siskind LJ, Kolesnick RN, Colombini M (2002) Ceramide channels increase the permeability of the mitochondrial outer membrane to small proteins. J Biol Chem 277: 26796–26803

Stahl E, Hartmann M, Scholten N, Zeier J (2019) A Role for tocopherol biosynthesis in Arabidopsis basal immunity to bacterial infection. Plant Physiol 181: 1008–1028

Stahl E, Brillatz T, Queiroz EF, Marcourt L, Schmiesing A, Hilfiker O, Riezman I, Riezman H, Wolfender JL, Reymond P (2020) Phosphatidylcholines from pieris brassicae eggs activate an immune response in arabidopsis. eLife 9: e60293.

Stotz HU, Mueller S, Zoeller M, Mueller J, Berger, S (2013) TGA transcription factors and jasmonate-independent COI1 signalling regulate specific plant responses to reactive oxylipins. J Exp Bot 64: 963–975

Stuart J (2015) Insect effectors and gene-for-gene interactions with host plants. Curr Opin Insect Sci 9: 56–61

Suzuki Y, Sogawa K, Seino Y (1996) Ovicidal reaction of rice plants against the whitebacked planthopper, *Sogatella furcifera* HORVATH (Homoptera: Delphacidae). Appl Entomol Zool 31: 111–118

Ternes P, Feussner K, Werner S, Lerche J, Iven T, Heilmann I, Riezman H, Feussner I (2011) Disruption of the ceramide synthase LOH1 causes spontaneous cell death in *Arabidopsis thaliana*. New Phytol 192: 841–854

Townley HE, McDonald K, Jenkins GI, Knight MR, Leaver CJ (2005) Ceramides induce programmed cell death in Arabidopsis cells in a calcium-dependent manner. Biol Chem 386: 161–166

Wang W, Yang X, Tangchaiburana S, Ndeh R, Markham JE, Tsegaye Y, Dunn TM, Wang GL, Bellizzi M, Parsons JF (2008) An inositolphosphorylceramide synthase is involved in regulation of plant programmed cell death associated with defense in Arabidopsis. Plant Cell 20: 3163–79

Weber H, Chételat A, Reymond P, Farmer EE (2004) Selective and powerful stress gene expression in Arabidopsis in response to malondialdehyde. Plant J 37: 877–888

Wright C, Beattie G (2004) *Pseudomonas syringae* pv. tomato cells encounter inhibitory levels of water stress during the hypersensitive response of *Arabidopsis thaliana*. Proc Natl Aca Sci USA 101: 3269–3274

Wu JX, Li J, Liu Z, Yin J, Chang ZY, Rong C, Wu JL, Bi FC, Yao N (2015) The Arabidopsis ceramidase AtACER functions in disease resistance and salt tolerance. Plant J 81: 767–780

Yamasaki M, Yoshimura A, Yasui H (2003) Genetic basis of ovicidal response to whitebacked planthopper (*Sogatella furcifera* Horváth) in rice (*Oryza sativa* L.). Mol Breed 12: 133–143

Yang Y, Xu J, Leng Y, Xiong G, Hu J, Zhang G, Huang L, Wang L, Guo L, Li J, et al. (2014) Quantitative trait loci identification, fine mapping and gene expression profiling for ovicidal response to whitebacked planthopper (*Sogatella furcifera* Horvath) in rice (*Oryza sativa* L.). BMC Plant Biol 14: 145

Young M, Kester, M, Wang H-G (2013) Sphingolipids: regulators of crosstalk between apoptosis and autophagy. J Lipid Res 54: 5–19

Zvereva AS, Golyaev V, Turco S, Gubaeva EG, Rajeswaran R, Schepetilnikov MV, Srour O, Ryabova LA, Boller T, Pooggin MM (2016) Viral protein suppresses oxidative burst and salicylic acid-dependent autophagy and facilitates bacterial growth on virus-infected plants. New Phytol 211: 1020–1034

